# Aberrant extracellular matrix and cardiac development in models lacking the PR-DUB component ASXL3

**DOI:** 10.1101/2022.07.14.500124

**Authors:** BT McGrath, YC Tsan, S Salvi, N Ghali, DM Martin, M Hannibal, CE Keegan, A Helms, A Srivastava, SL Bielas

## Abstract

**Background:** Clinical and research based genetic testing has uncovered genes that encode chromatin modifying complex components required for organogenesis. Covalent histone modifications play a key role in establishing transcriptional plasticity during development, required for cell fate specification, and have been implicated as a developmental mechanism that accounts for autism spectrum disorder (ASD) and CHD co-occurrence. *ASXL3* has been identified as a high confidence ASD gene. ASXL3 is a component of the Polycomb Repressive Deubiquitination (PR-DUB) complex, which deubiquitinates histone H2A. However, the role of ASXL3 in cardiac development remains unknown.

**Methods:** We used CRISPR/Cas9 gene editing to generate clinically relevant *Asxl3* frameshift alleles in a mouse model and human embryonic stem cells (hESCs). To evaluate ASXL3 function in developing hearts, we performed structural, molecular, immunostaining and histological analyses. Transcriptomic and cellular compositional changes were assessed with bulk RNA sequencing of mouse hearts and single-cell RNA sequencing (scRNA-seq) of human cardiac tissue differentiated from isogenic hESC lines.

**Results:** Biallelic genetic inactivation of *Asxl3* leads to perinatal lethality and increased levels of histone H2A mono-ubiquitination, which are regulated by PR-DUB. *Asxl3^+/fs^* and *Asxl3^fs/fs^* mice display cardiac abnormalities including ventricular hypoplasia, septal defects, and bifid cardiac apex with variable penetrance. The presence of underdeveloped ventricles is preceded by increased progenitor proliferation in the ventricles, as determined by EdU incorporation. Differential gene expression, assessed by bulk RNA sequencing implicates extracellular matrix dysfunction as a pathogenic mechanism. This correlates with a reduction in vimentin-positive cardiac fibroblasts. scRNA-seq of cardiac cultures differentiated from human *ASXL3^fs/fs^* ESC lines exhibit altered ratios of cardiac fibroblasts and cardiomyocytes. Similar to the mouse data, genes essential for extracellular matrix composition and signaling are differentially expressed between *ASXL3^+/+^* and *ASXL3^fs/fs^* human *in vitro* differentiated cardiac tissue. The observed transcriptomic changes predict diminished cell-cell signaling interactions between cardiac fibroblasts and cardiomyocyte progenitors in *ASXL3* cultures.

**Conclusions:** Collectively, our data implicates species-specific roles for ASXL3 in both human and mouse cardiac development. These results highlight the role of extracellular matrix gene programs by cardiac fibroblast during cardiomyocyte development and provide insight into mechanisms of altered cardiogenesis by autism risk genes.

## INTRODUCTION

Heart development requires exquisite orchestration of cardiac progenitor proliferation, migration, differentiation, morphogenesis and developmental remodeling to form four chambers. Disruption of this complex process is the basis of congenital heart defects, the most prevalent birth defect. Investigating the genetic and molecular mechanisms underlying cardiac development will unveil the transcriptomic and regulatory networks essential for normal and abnormal cardiogenesis.

Transformation of the embryonic heart tube into the mammalian four chambered heart requires coordinated proliferation of cardiac progenitor cell (CPC) pools and differentiation toward diverse cardiac lineages. Analysis of cardiogenesis in model organisms has allowed the developmental origin of the mature structural heart features to be traced back to one of the two mesoderm derived cardiac primordia, the first (FHF) and second heart fields (SHF), or neural crest lineages [1]. The left ventricle and atria are a product of FHF progenitors and the SHF contributes to the right ventricle, atria, outflow and inflow tracks. Single-cell transcriptomics studies along heart organogenesis have defined the shared and distinct expression profile of CPCs progressing through a series of intermediate cardiac-cell states to mature endocardial and cardiomyocyte cardiac lineages, in mouse and human model systems [2–5].

Within the embryonic heart, the extracellular matrix (ECM) plays an active role in governing the spatial programs of cardiac organogenesis[6]. Beyond structural support for cardiac cells, the ECM influences cardiomyocyte migration during cardiac tube expansion, looping, ventricular trabeculation and compaction and regulation of cell proliferation and differentiation [7–16]. The influence of the ECM on these developmental processes depends on the coordinated synthesis and degradation of ECM secreted factors and expression of corresponding ECM receptors [6]. For instance, cardiac fibroblasts, which comprise a large cell population in the heart, secrete high levels of developmental stage-specific extracellular matrix and growth factors, including fibronectin (FN) [17].Cell surface expression of β1 Integrin by cardiomyocytes binds FN in the ECM, promoting CPC proliferation and expansion of ventricular chambers during cardiogenesis [14, 18, 19]. ECM composition varies across cardiac structures and development. Independent of the stage of heart development, ECM composition is transcriptionally regulated across various cardiac cell lineages and can be disrupted by epigenetic mechanisms that alter chromatin remodeling or genome wide post-translational modifications[12, 20, 21].

During cardiogenesis, epigenetic mechanisms establish transcriptional plasticity that generates distinct transcriptional profiles for development. The evolutionarily conserved Polycomb group (PcG) complexes contribute to establishing this transcriptional plasticity by regulating transcriptional repression [22]. The PcG complexes, Polycomb repressive complex 1 (PRC1), complex 2 (PRC2) and PR-DUB, are subdivided on the basis of associated enzymatic activities[23]. Histone H3 lysine 27 trimethylation (H3K27me3) is catalyzed by PRC2. PRC1 catalyzes mono-ubiquitylation of H2A (H2AUb1), while PR-DUB hydrolyzes H2AUb1 deubiquitination. The core components of the PR-DUB complex are the ubiquitin hydrolase BAP1 (BRCA1 associated protein 1) and one of three ASXL family members, ASXL1, ASXL2 or ASXL3 [24]. Individual ASXL family members function as the mutually exclusive obligate regulatory subunits of the PR-DUB complex, as BAP1 does not exhibit histone H2A deubiquitination activity (DUB) in the absence of an ASXL protein [25, 26]. While all ASXL family members were expressed in the embryonic mouse heart, the role of Asxl3 during cardiac development has not been investigated [27].

Monoallelic frameshift *ASXL3* variants are the genetic basis of Bainbridge-Ropers Syndrome (BRS) and syndromic autism spectrum disorder (ASD), characterized by repetitive and social features of ASD, global developmental delay, intellectual disability, hypotonia, craniofacial dysmorphism, reduced gastrointestinal motility and altered sleep cycle [28, 29]. While congenital heart defects are not enriched in this cohort, they have been described as a feature of ultra-rare biallelic recessive *ASXL3* missense variants [30]. We investigated the role of ASXL3 in cardiac development using biallelic frameshift *Asxl3* mouse model and cardiac tissue differentiated from biallelic frameshift *ASXL3* human embryonic stem cell (hESC) lines. Constitutive loss of *Asxl3* exhibit highly penetrant CHDs and perinatal lethality, with less severe cardiac phenotypes observed in heterozygous animals. Altered composition of cardiac cell lineages and ECM components were observed in models from both species. Transcriptomics implicate species- specific genetic and molecular features of these shared pathogenic mechanisms. Correlation between these biallelic models highlight a dosage-sensitive role for *ASXL3* in cardiogenesis and underscores the differential expressivity of CHDs in BRS/ASD.

## RESULTS

### H2AUb1 elevated in Asxl3 frameshift mouse model

To investigate the role of ASXL3 in development, we generated a mouse model with a two base-pair deletion in *Asxl3*, corresponding to homologous nucleotides classified as pathogenic *ASXL3* BRS/ASD variants. Single guide RNAs (sgRNAs) were designed to target a region of mouse exon 12, which is homologous to human exon 11 (Figure 1A). CRISPR/Cas9 editing created an *Asxl3* c.990_992delCA:993T > G;p.T484Afs*5 frameshift (*Asxl3fs*) allele through non- homologous end joining in B6SJL hybrid blastocyst after pronuclear injection (Figure 1B). Expression of full-length *Asxl3* and its protein product was not detected by RT-qPCR and Western blot in *Asxl3^fs/fs^* mice (Figure 1C and 1D). ASXL3 interacts with BAP1 to form the PR-DUB complex, which mediates histone H2A deubiquitylation [29]. We measured the levels of Histone H2A mono-ubiquitination in E13.5 hearts by Western blot. Consistent with findings from patient derived fibroblasts, we found a 1.5 fold increase of H2AUb1 relative to histone H3 in *Asxl3^fs/fs^* mice (Figure 1E and 1F).

**Figure 1.**
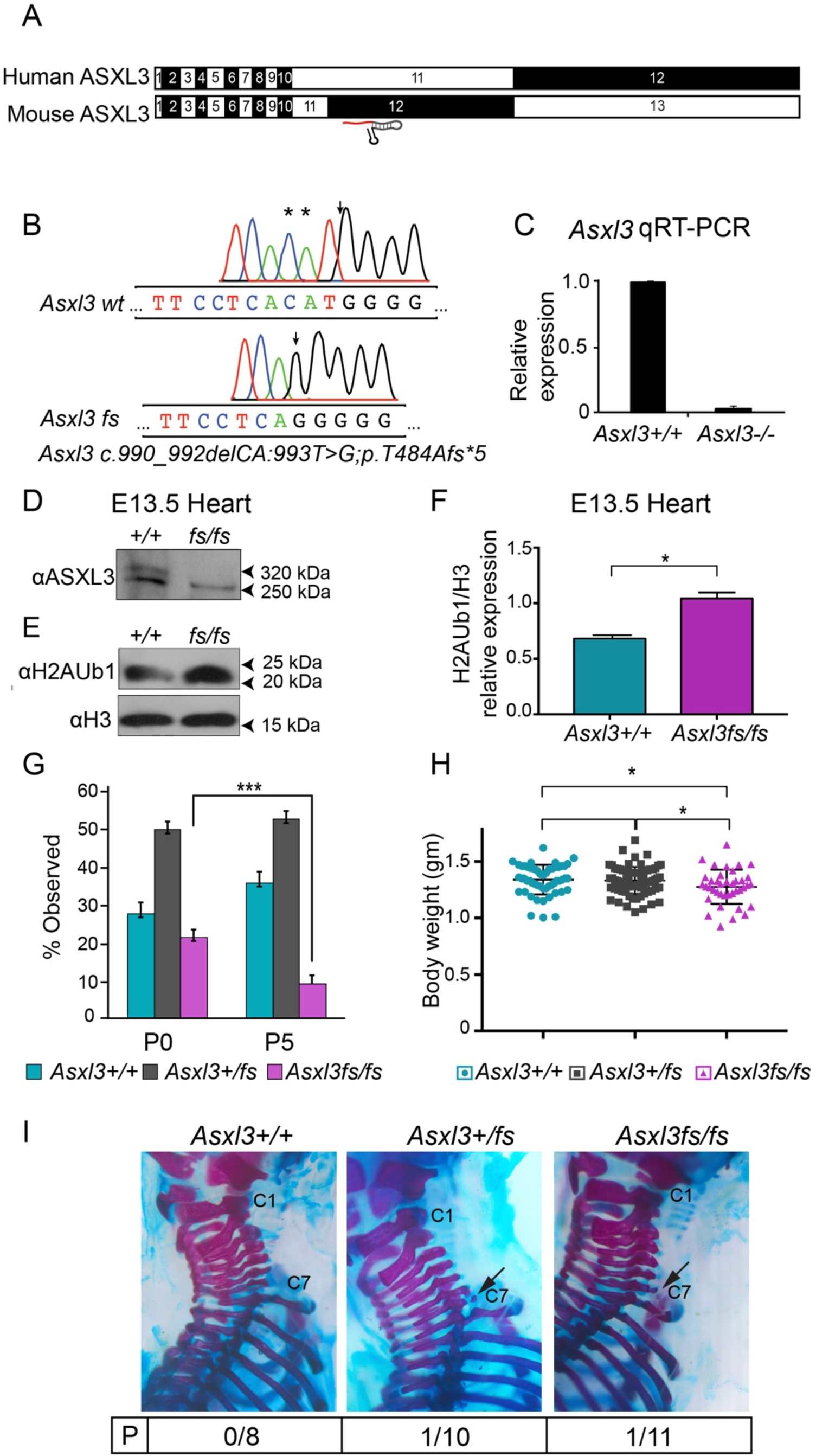
Generation of *Asxl3^fs/fs^* mice. **A,** Depiction of CRISPR/Cas9 genome editing strategy for *Asxl3^fs^* mouse generation. The sgRNA targets a loci of mouse *Asxl3* exon 12 that is homologous to a region of human *ASXL3* exon 11 that contains clinically relevant variants. **B,** Representative indel sequences from Sanger sequencing of targeted *Asxl3* loci. **C,** RT-qPCR analysis of Asxl3 mRNA levels in Embryonic day 13.5 (E13.5) *Asxl3^fs^* edited neural progenitor cells. **D,** Western blot analysis shows the expression of full length ASXL3 is lost in E13.5 *Asxl3^fs/fs^* hearts. **E,** Western blot analysis of H2AK119Ub1 (H2AUb1) and H3 acid extracted histones from *Asxl3^+/+^* and *Asxl3^fs/fs^* E13.5 hearts. **F,** Quantification of H2AUb1 relative to H3 control levels revealed a 2-fold increase in *Asxl3^fs/fs^* E13.5 hearts compared to control. **G,** Percentage of observed *Asxl3^+/+^, Asxl3^+/fs^,* and *Asxl3^fs/fs^* mice at P0 and P5. Chi square analysis showed a reduction in *Asxl3^fs/fs^* mice based on normal mendelian distribution. **H,** Student’s *t*-test shows a significant reduction in body weight of *Asxl3^fs/fs^* P0 mice relative to wild type and heterozygous mice (n=53 for *Asxl3^+/+^*, n=75 for *Asxl3^+/fs^* and n=39 for *Asxl3^fs/fs^*). **I,** Alcian blue and Alizarin red staining of P0.5 embryos. Arrows indicate cervical ribs observed in *Asxl3^+/fs^,* and *Asxl3^fs/fs^* mice. Penetrance of cervical ribs by genotype displayed below. Values are mean±SEM. \**P*<0.05, \*\**P*<0.01, \*\*\**P*<0.001.

### Asxl3 null mouse exhibits Polycomb transcriptional repression phenotypes

Heterozygous intercrosses yielded *Asxl3^fs/fs^* at standard Mendelian ratios till birth at P0.5 (Figure 1G). Consistent with neonatal lethality, *Asxl3^fs/fs^* pups are underrepresented from P0.5 to P5. At birth, *Asxl3^fs/fs^* mice weigh less than both *Asxl3^+/fs^* and *Asxl3^+/+^* littermates which are comparable in weight (Figure 1H). Given the important role for Polycomb transcriptional repression on Hox gene expression, we performed skeletal alcian blue staining to assess evidence of skeletal homeotic transformations [23]. No defects were observed in the vertebral column of 8 *Asxl3^+/+^* pups, while a low penetrant cervical rib homeotic transformation of cervical vertebrae C7 was observed for 1 in 10 *Asxl3^+/fs^,* and 1 of 11 *Asxl3^fs/fs^* pups. Small ossified rib anlagen at cervical vertebra C7 are observed at very low penetrance in wild-type mice of some hybrid mouse strains [31]. This phenotype has not been described for a B6SJL hybrid mouse strain, implicating a nonredundant role of ASXL3 in Hox gene regulation of axial skeleton segmentation.

### Asxl3 mutant neonates exhibit severe congenital heart defects

Phenotypic evaluation of multiple organ systems was performed to determine an etiology of neonatal lethality. Stomach milk is externally observed in P0.5 *Asxl3^fs/fs^* pups, suggesting that suckling reflexes are intact. Hematoxylin and eosin (H&E) histological evaluation of whole mount P0.5 *Asxl3^+/+^*, *Asxl3^+/fs^* and *Asxl3^fs/fs^* neonates depicted normal gross anatomy of liver, lungs, kidneys, small intestine, colon, testis, and stomach for all three genotypes (Figure 1 in the Data Supplement). No structural brain defects were observed by cresyl violet staining, but distinct borders defining cortical layers are diminished (data not shown). These defects are not predicted to result in neonatal lethality.

Conversely, *Asxl3^fs/fs^* neonates exhibit congenital cardiac defects by H&E staining and gross organ structural defects. Hearts of P0.5 mice displayed congenital morphological abnormalities ranging from cardiac bifida to separated ventricles in *Asxl3^+/fs^* and *Asxl3^fs/fs^* mice at increasing penetrance, respectively (Figure 2A and 2B). Bifida cardiac apex was observed in 16.7% of *Asxl3^fs/fs^* mice and separated ventricles were observed in 4.5% of *Asxl3^+/fs^* and 16.7% of *Asxl3^fs/fs^* mice. Out of 39 *Asxl3^fs/fs^* neonatal hearts serially sectioned and analyzed by H&E staining, 11 (28%) showed right ventricular hypoplasia, with 4 presenting with almost complete loss of the ventricular lumen, making this the most penetrant heart phenotype in this *Asxl3* mouse model. Apical muscular ventricular septal defects were also detected in this analysis in 5.1% of *Asxl3^fs/fs^* mice. The ventricles in hypoplastic ventricular hearts appeared incomplete with reduced luminal space throughout the entirety of serial sections (Figure 2 in the Data Supplement). Such cardiac defects have been observed in transgenic mouse models of genes that encode components of Polycomb complexes and are predicted to be the basis of neonatal lethality [27].

**Figure 2.**
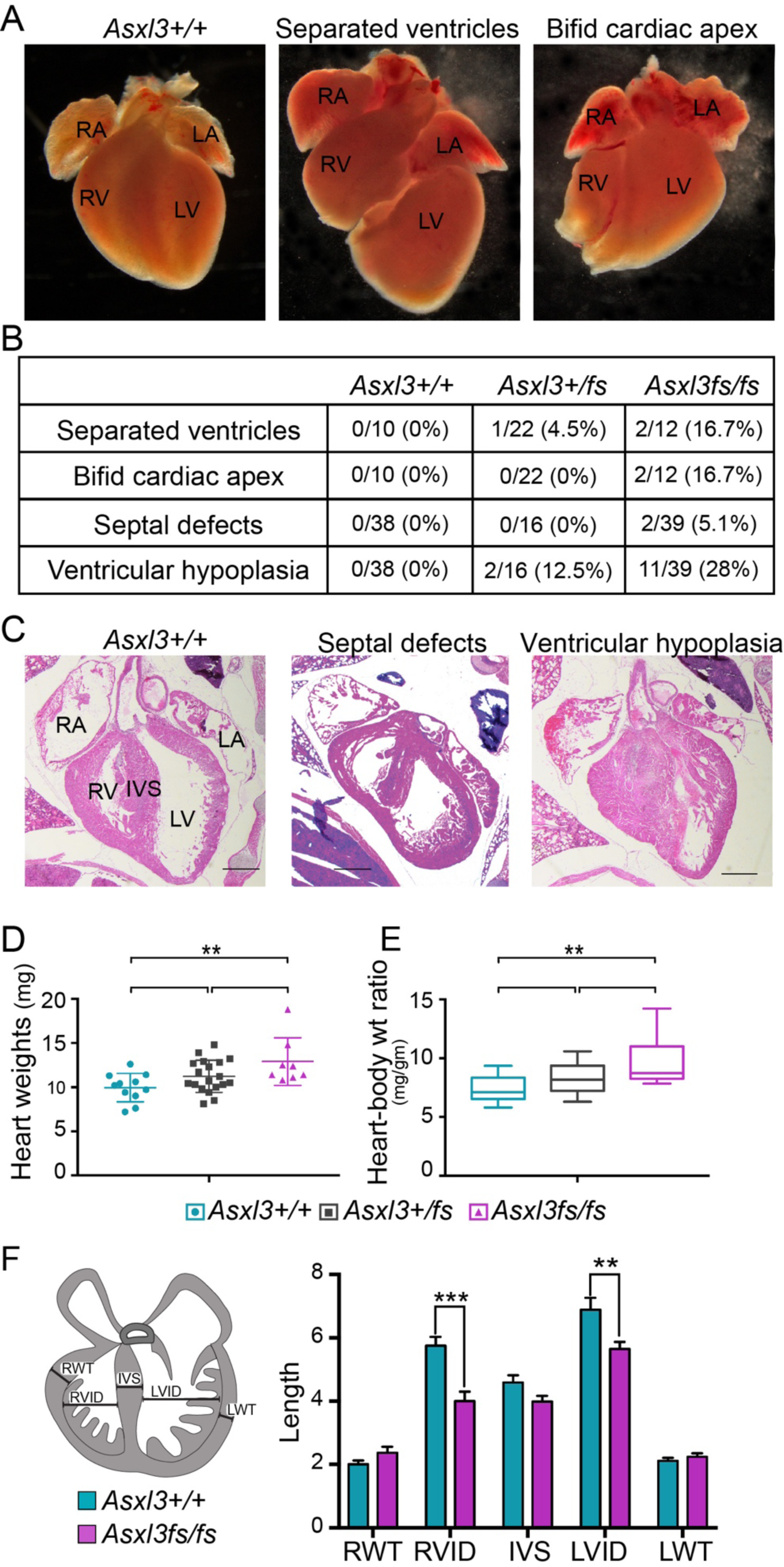
Representative *Asxl3^fs^*-associated cardiac anomalies. **A,** Whole mount images of P0 dissected hearts displaying separated ventricles and bifid cardiac apex. **B,** Table summarizing cardiac anomalies observed in *Asxl3^+/+^, Asxl3^+/fs^,* and *Asxl3^fs/fs^* P0 mice. **C,** Hematoxylin and eosin staining of P0 hearts shows representative images of the *Asxl3^fs^*-associated septal defects and ventricular hypoplasia. Scale bars, 500 µm. Student’s *t-*test analysis of P0 **D,** heart weights and **E,** heart to body weight ratio between genotypes (n=11 for *Asxl3^+/+^*, n=18 for *Asxl3^+/fs^* and n=8 for *Asxl3^fs/fs^*). Data are presented as mean±SEM. **F,** P0 heart Illustration detailing morphometric measurements for right wall thickness (RWT), right ventricular free wall (RVFW), interventricular septum (IVS), left ventricular free wall (LVFW), left wall thickness (LWT). **G,** Student’s *t*-test analysis of P0 morphometric measurements shows increased RVFW and LVFW in *Asxl3^fs/fs^* mice compared to *Asxl3^+/+^* (n=4 for *Asxl3^+/+^*, n=5 for *Asxl3^fs/fs^*). \**P*<0.05, \*\**P*<0.01, \*\*\**P*<0.001.

Right ventricular hypoplasia in *Asxl3^+/fs^* and *Asxl3^fs/fs^* P0.5 hearts correlates to increased heart weight and heart to body weight ratios, consistent with increased cardiac tissue. We independently weighed P0.5 pups as well as dissected hearts with extra-cardiac tissue removed. *Asxl3^fs/fs^* hearts weigh significantly more than *Asxl3^+/+^* hearts, with a corresponding increase in heart to body-weight ratio (Figure 2D and 2E). No significant difference was observed in heterozygous mice. Quantitative morphometric analysis of H&E stained heart sections were performed to characterize the ventricular tissue changes in hypoplastic hearts. Morphometric analysis included interventricular septum (IVS), left ventricle internal diameter (LVID), left wall thickness (LWT), right ventricular internal diameter (RVID), and right wall thickness (RWT) (Figure 2F). This analysis revealed a decrease in LVID and RVID in *Asxl3^fs/fs^* mice consistent with ventricular hypoplasia (Figure 2G). The absence of a significant increase in RWT, LWT, or IVS length could be due to the variable penetrance of hypoplastic hearts since the septum and ventricular walls of hypoplastic hearts appear thicker (Figure 1 in the Data Supplement).

### Increased proliferation during heart development

Several pathogenic mechanisms of cardiac development including fibrosis, increased cardiomyocyte size (hypertrophy) or altered cardiac progenitor proliferation (hyperplasia) could account for an increased ventricular chamber wall thickness. Masson trichrome staining performed on P0.5 paraffin-embedded heart sections did not show evidence of fibrosis in ventricular or septum tissue from mice of any genotype (Figure 3A and 3B). To investigate the possibility of cardiomyocyte hypertrophy, the cross-sectional area of cardiomyocytes was quantified from P0.5 coronal cardiac sections stained with Alexa 488 conjugated wheat germ agglutinin (WGA). No significant difference in cardiomyocyte size was observed in *Asxl3^+/fs^* or *Asxl3^fs/fs^* tissue relative to controls across corresponding structural areas (Figure 3 in Data Supplement).

**Figure 3.**
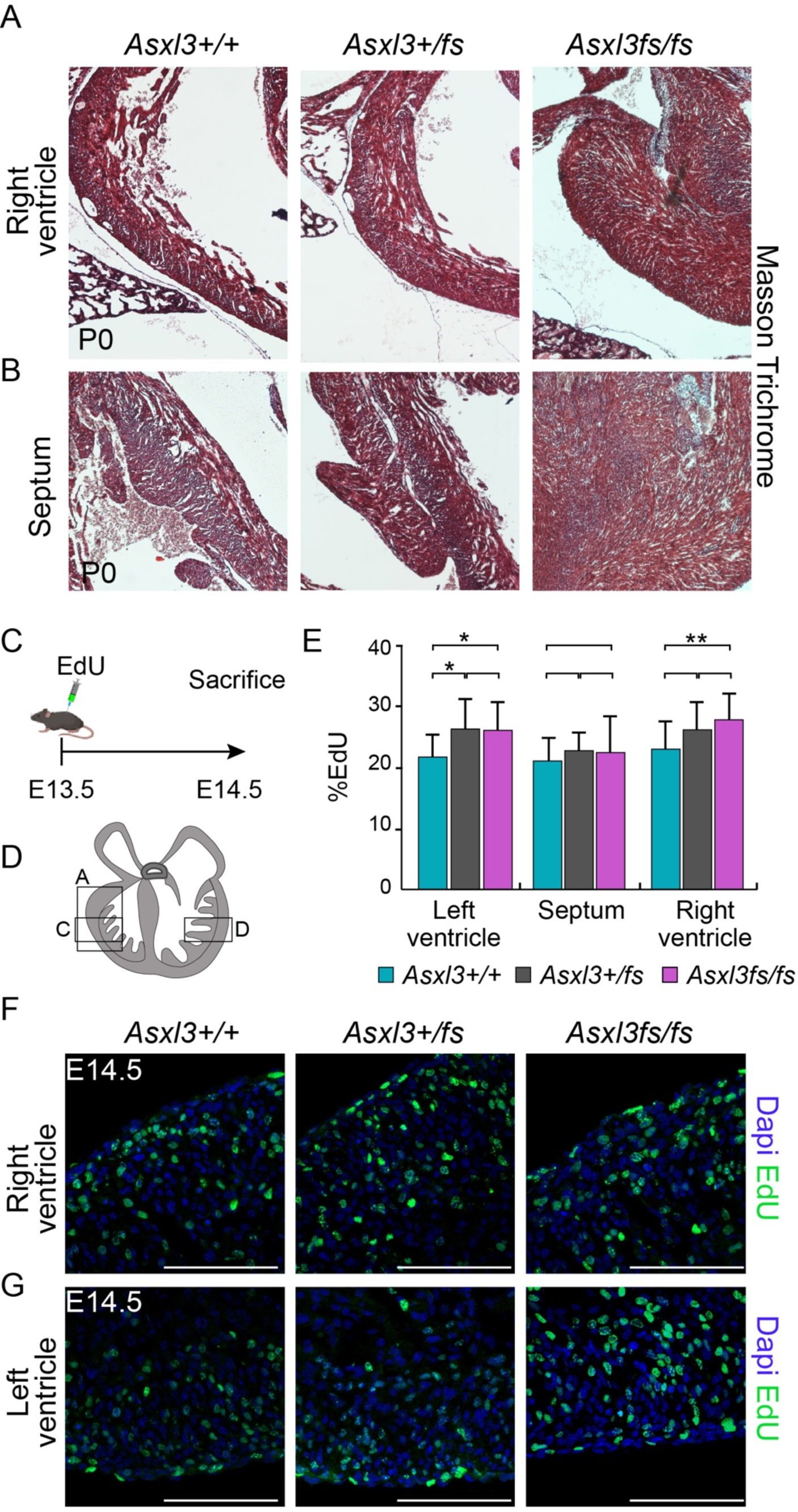
Increased proliferation during heart development. Representative images of the **A,** right ventricular walls and **B,** septums from *Asxl3^+/+^, Asxl3^+/fs^,* and *Asxl3^fs/fs^* P0 hearts stained with Masson trichrome. Scale bars, µm. No difference in Masson trichrome staining was observed between genotypes. **C,** Experimental timeline showing EdU labeling at E13.5 and collection 24 hours later. **D,** Schematic depicting representative regions of images used for quantifications in transverse heart section **E,** Student’s *t-*test analysis of EdU-positive proliferative cells in E14 hearts between genotypes (n=4 for *Asxl3^+/^*^+^, n=10 for *Asxl3^+/fs^* and n=5 for *Asxl3^fs/fs^*). Representative images of E14.5 **F,** right and **G,** left ventricles from *Asxl3^+/+^*, *Asxl3^+/fs^* and *Asxl3^fs/fs^* hearts after EdU incorporation. Scale bars, 100 µm. Values are Values are mean±SEM. \**P*<0.05, \*\**P*<0.01, \*\*\**P*<0.001.

CPC proliferation during development was interrogated using a 24hr pulse of the EdU, to label mitotically active undergoing DNA replication. Timed-pregnant dames were administered EdU, by intraperitoneal injection at embryonic day 13 (E13) (Figure 3C). A day later, hearts from E14 littermates were fixed, cryosectioned and EdU was detected through a Click-it reaction (Figure 3F and 3G). No appreciable changes in EdU positive cells were noted in the septum between genotypes (Figure 3E). In the left ventricle of *Asxl3^+/fs^* and *Asxl3^fs/fs^* hearts, EdU was detected in more cells relative to wild type littermates (Figure 3E). EdU-positive cells were also increased in the right ventricle of *Asxl3^+/fs^* and *Asxl3^fs/fs^* hearts, however, significance was achieved in only *Asxl3^fs/fs^* tissue relative to controls (Figure 3E). Overall, *Asxl3^fs/fs^* hearts tend to show increased levels of EdU incorporation in both the left and right ventricles at E14. This is consistent with a model of cardiac progenitor hyperplasia, with a secondary reduction in ventricular luminal size.

### RNA Seq uncovers distinct gene expressions in E18.5 Asxl3 mutants

*De novo* and CRISPR-edited *ASXL3* genetic variants disrupt ASXL3-dependent PR-DUB H2AUb1 deubiquitination and transcriptional regulation (Figure 1E). To analyze transcriptional changes correlated with elevated H2AUb1 in *Asxl3^+/fs^* and *Asxl3^fs/fs^* hearts, we performed bulk- RNA sequencing of E18.5 ventricles isolated from closely associated aorta, left atrium and right atrium from four sets of *Asxl3-*littermates. Highly reproducible transcriptomic changes were observed in the ventricular samples from both *Asxl3^+/fs^* and *Asxl3^fs/fs^* samples. 333 differentially expressed genes (DEGs; false discovery rate adjusted *P*-values defined by DESeq2, *q* < 0.05 and log2 fold change of ≥0.4) were identified in *Asxl3^+/fs^* ventricles, with 170 (51.05%) downregulated and 163 (48.95%) upregulated (Figure 4A). Similarly, 366 DEGs were observed in *Asxl3^fs/fs^* ventricular tissue with 161 (43.99%) downregulated and 205 (56.01%) upregulated (Figure 4B).

**Figure 4.**
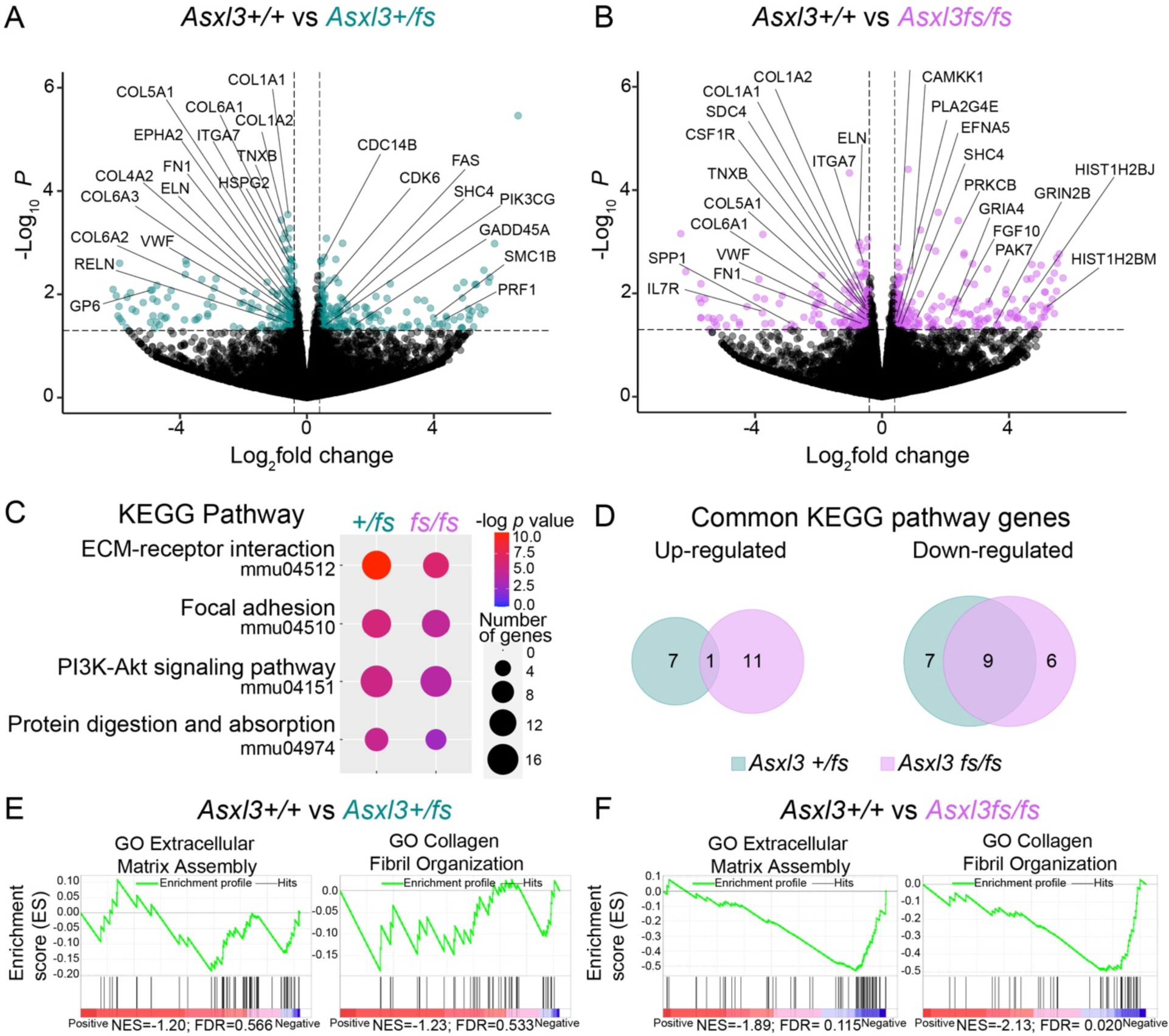
Loss of Asxl3 leads to altered expression of extracellular matrix components. Volcano plots exhibiting gene expression changes in **A,** *Asxl3^+/fs^* vs. *Asxl3^+/+^* and **B,** *Asxl3^fs/fs^* vs. *Asxl3^+/+^* E18.5 hearts. Genes with a p-value > 0.05 and log fold change > 0.4 are highlighted. GSEA enrichment plots for GO:Extracellular Matrix Assembly and GO:Collagen Fibril Organization from **C,** *Asxl3^+/fs^* vs. *Asxl3^+/+^* and **D,** *Asxl3^fs/fs^* vs. *Asxl3^+/+^* analysis. Gene set enrichment analysis (GSEA) was performed with the ontology gene sets in GSEA molecular signature database. **E,** KEGG pathway enrichment analysis for *Asxl3^+/fs^* and *Asxl3^fs/fs^* downregulated genes. The top 4 KEGG pathways for both *Asxl3^+/fs^* and *Asxl3^fs/fs^* are shown. The size of each dot represents the gene number and the shading represents the -log10 p-value. **F,** The Venn diagram illustrates the overlap of unique upregulated (left) or downregulated (right) genes contributing to common KEGG pathways.

Comparison of all *Asxl3^+/fs^* and *Asxl3^fs/fs^* DEGs revealed that ∼30% of DEGs were represented in both genotypes (Figure 4A and 4B in the Data Supplement). Kyoto Encyclopedia of Genes and Genomes (KEGG) pathway enrichment analysis of *Asxl3^+/fs^* and *Asxl3^fs/fs^* transcriptomics revealed overlapping pathways for downregulated DEGs including ‘ECM-receptor interactions’, ‘focal adhesion’, and ‘PI3K-AKT signaling pathway’ (Figure 4C). Similar findings were detected with GO analysis (Figure 4C in Data Supplement). Together these altered pathways implicate disrupted extracellular matrix (ECM) function in *Asxl3^+/fs^* and *Asxl3^fs/fs^* ventricles. The ECM provides both signaling and structural functions in the developing heart that are critical for the heart organogenesis [6]. Downregulated DEGs enriched in *Asxl3^+/fs^* and *Asxl3^fs/fs^* shared pathways include important cardiac ECM components like *Col1a1, Col5a1, Fbln2, Fbn1, Eln, Ltbp4, and Fn1*, (Figure 4A and 4B). Of note, while *Asxl3^+/fs^* and *Asxl3^fs/fs^* shared a similar number of upregulated (55) and downregulated (52) genes, upregulated DEGs enriched in KEGG pathways were not shared between *Asxl3^+/fs^* and *Asxl3^fs/fs^* datasets (Figure 4D). ECM involvement in *Asxl3^fs/fs^* cardiac phenotype was also implicated by gene set enrichment analysis (GSEA) of the C5 ontology gene set from the molecular signature database (MSigDB), which also identified significant DEG transcript enrichment in GO Extracellular Matrix Assembly and GO Collagen Fibril Organization (Figure 4E and 4F). While negative normalized enrichment scores (NESs) quantified the correlation of downregulated DEGs for both *Asxl3^+/fs^* and *Asxl3^fs/fs^* samples, this analysis also reveals the intermediate transcriptional changes for *Asxl3^+/fs^* relative to *Asxl3^fs/fs^* samples, for both the rank order DEGs and negative NES. An *Asxl3^+/fs^* NES of -1.2 and -1.23 versus *Asxl3^fs/fs^* NES of -1.89 and -2.13 for GO Extracellular Matrix Assembly and GO Collagen Fibril Organization pathways respectively quantify the difference of enriched hits from bottom-ranked genes in the pathway. This data does not differentiate between timing of the onset of differential expression across cardiac development, both of which may be required to describe the discrepancy in the severity of cardiac phenotypes between *Asxl3^+/fs^* and *Asxl3^fs/fs^* animals.

### Reduced cardiac fibroblasts in ventricles of *Asxl3* mutants

Cardiac fibroblasts are a prominent source of ECM deposition, degradation, and remodeling in the developing heart [17]. Embryonic cardiac fibroblasts are also a source of integrin ligands, including fibronectin and collagen, that promote cardiomyocyte proliferation through integrin signaling pathways [14]. Given the differential expression of ECM components we detected in *Asxl3^+/fs^* and *Asxl3^fs/fs^* ventricles and increased proliferation observed at E14, cardiac fibroblasts distribution and ECM deposition was evaluated. Coronal sections of fixed P0.5 heart were immunostained for vimentin, an intermediate filament protein marker of cardiac fibroblasts. Vimentin-positive cells were quantified in each ventricle separately and the septum (Figure 5C). An equivalent number of fibroblasts were observed in the septum between all three genotypes (Figure 5B). A significant reduction in vimentin-positive cells were quantified in *Asxl3^+/fs^* and *Asxl3^fs/fs^* samples relative to controls (Figure 5D and 5F). This decrease in cardiac fibroblasts is a mechanism that would account for downregulation of ECM gene signatures by RNA sequencing.

**Figure 5.**
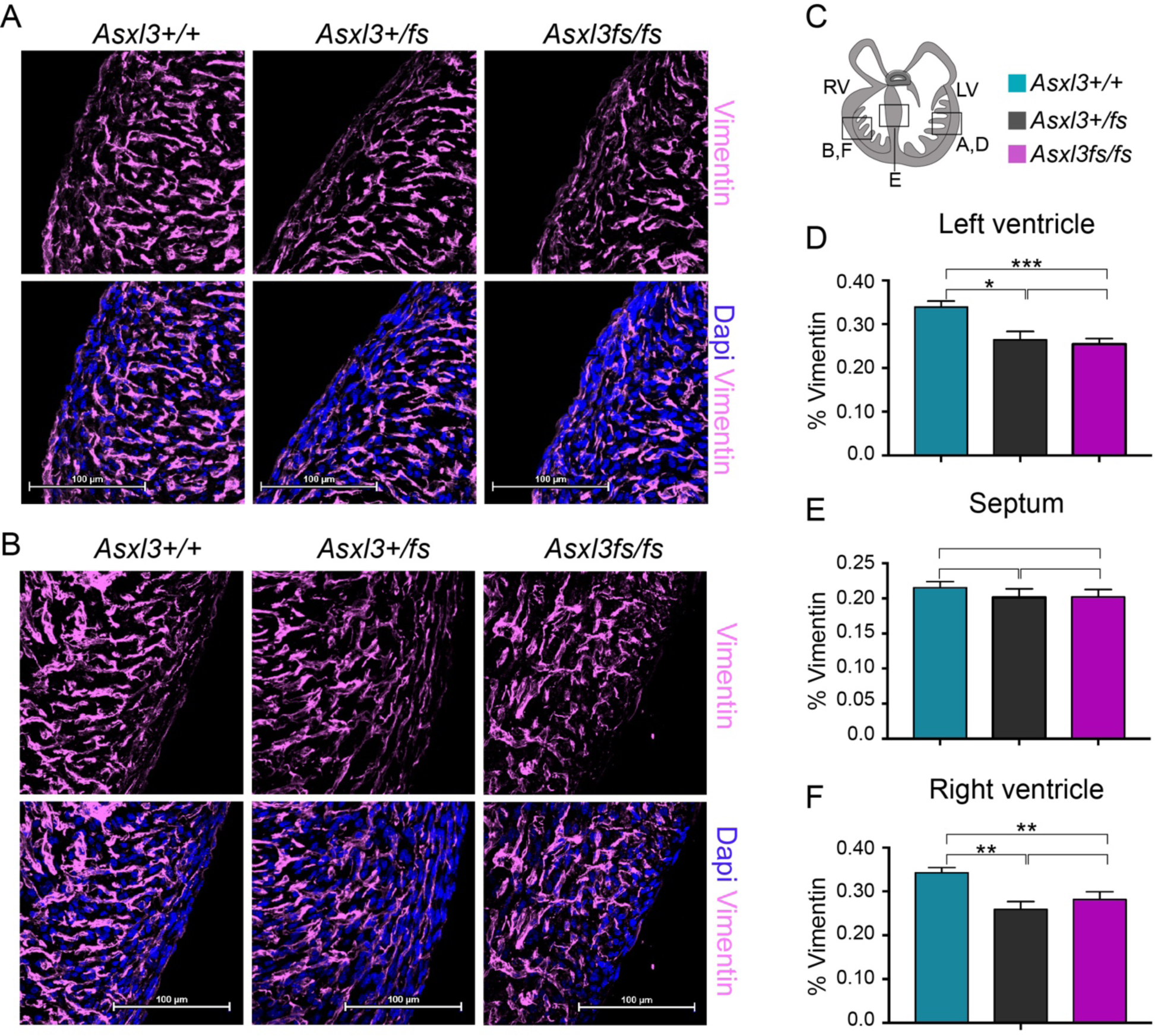
Reduction in vimentin-positive cardiac fibroblasts. Immunostaining of P0 **A,** left ventricular free wall and **B,** right ventricular free wall with vimentin to mark cardiac fibroblasts and nuclei stained with DAPI (n=6 for *Asxl3^+/+^*, n=3 for *Asxl3^+/fs^* and n=5 for *Asxl3^fs/fs^*). **C,** Schematic of P0 transverse heart sections depicting representative regions of images used for quantifications. Statistical analysis of vimentin-positive cells detected in the **D,** left ventricular free wall, **E,** interventricular septum, and **F,** right ventricular free wall of *Asxl3^+/+^*, *Asxl3^+/fs^* and *Asxl3^fs/fs^* P0 hearts. Values are mean ± SEM. RV indicates right ventricle; LV, left ventricle. \**P*<0.05, \*\**P*<0.01, \*\*\**P*<0.001.

*Collagen1A1* (*COL1A1*) and *Collagen3A1* (*Col3A1*) expression was downregulated by RNA-seq and ECM deposition in E15.5 and P0.5 heart were validated by immunohistochemistry. Surprisingly, minimal collagen differences were observed at P0.5 in any heart region (Figure 5 in Data Supplement). At E15 *Col3A1* appeared comparable in the left ventricle of Asxl3+/+ and *Asxl3^fs/fs^* mice, but it appeared slightly increased in the right ventricle of *Asxl3^fs/fs^* littermates. Evaluation of other ECM components and/or additional developmental time points would provide greater insights into the disrupted ECM dynamics. Together these findings suggest that inactivation of *Asxl3* in cardiac development may have downstream consequences such as the disrupted collagen and decreased number of cardiac fibroblasts as is observed in the heterozygous and null hearts.

### ASXL3-dependent *in vitro* human cardiac differentiation

Given the *Asxl3^fs/fs^* mouse cardiac phenotype, and congenital heart defects attributed to biallelic *ASXL3* missense variants, a human *in vitro* model of cardiac differentiation was used to explore the function of *ASXL3* in normal physiology and pathology. CRISPR/cas9 gene-editing was used to create isogenic H9 human embryonic stem cell (hESC) lines carrying biallelic *ASXL3* c.1393dupT;p.C465LfsX4; and c.1390_1393del;p.E464AfsX19 (*ASXL3^fs/fs^*) variants in exon 11 an *ASXL3* exon enriched for variants clinically classified as pathogenic [29] (Fig 6A and Supp 6). Isogenic control and *ASXL3^fs/fs^* hESC lines were differentiated to a cardiac lineage using an established protocol, but lactate selection or magnetic beads assisted cell sorting was omitted to preserve non-cardiomyocyte lineages in the culture, thus enhancing the cellular heterogeneity of tissue differentiated from the common cardiogenic mesodermal lineage [32]. Differentiation of monolayer cultured hESC towards cardiomyocyte lineage is induced by modulating components of the WNT signaling pathway over first 5 successive days to establish cardiogenic mesoderm and transition into CPCs. By omitting cardiomyocyte selection at day 10, differentiation proceeds to generate NKX2-5 cardiomyocytes lineages and non-contractile cardiac derivatives (Figure 6B). Single-cell RNA sequencing was performed on *ASXL3^+/+^* control and *ASXL3^fs/fs^* cultures at 12 days of differentiations to characterize the progeny of cardiac lineages. A total of 18,577 cells (9,379 *ASXL3^+/+^* and 9,198 *ASXL3^fs/fs^* cells) with an average of 1001.73 genes detected per cell that met our data quality control and filtering parameters, were captured from three *ASXL3^+/+^* and two *ASXL3^fs/fs^* independent cardiac differentiations. The filtered dataset was analyzed using our pipeline based on Seurat version 3 [33].

**Figure 6.**
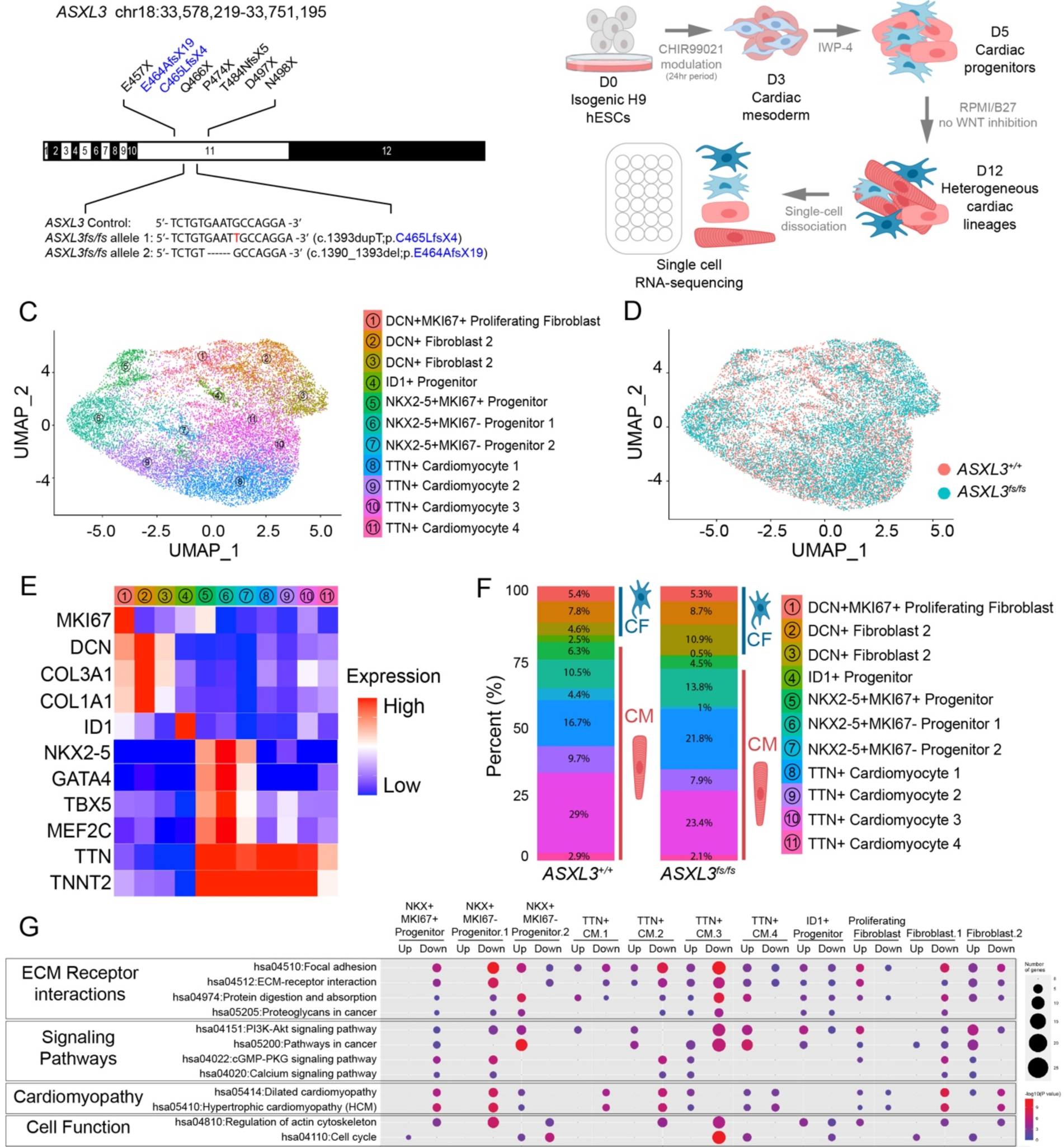
scRNA-seq analysis of *ASXL3^+/+^* and *ASXl3^fs/fs^* cardiac directed differentiation. **A**, *ASXL3* Frameshift mutations (blue) introduced in control H9 human embryonic stem cells via CRISPR gene editing and representative mutations identified in Bainbridge-Ropers Syndrome patients (black). The introduced frameshift mutations (*c.1393dupT; p.C465LfsX4* and *c.1390_1393del; p.E464AfsX19*) are within the same genomic region as the patients mutations **B**, Schematic representation of human embryonic stem cell cardiac directed differentiation. Differentiation day 12 cells were collected for single-cell RNA sequencing analysis. **C**, Uniform Manifold Approximation and Projection (UMAP) analysis of *ASXL3^+/+^* (n=9,379) and *ASXL3^fs/fs^* (n=9,198) cells across the cardiac lineage. 11 clusters were identified. **D**, UMAP from **C** colored by genotype. **E**, Heatmap of marker genes used for cell identity. **F**, Stacked barplot displaying the proportion of clustered cells detected from *ASXL3^+/+^* and *ASXL3^fs/fs^* cultures within each phenotypic clusters. **G**, Highlights of KEGG analysis of differentially expressed genes in each cluster in the human *ASXL3^fs/fs^* ESC *in vitro* cardiac differentiation.

By performing dimensionality reduction and unsupervised clustering with uniform manifold approximation and projection (UMAP) we identified 11 unique cell populations (Fig 6C and D).

Clusters were manually annotated according to expression of cell type markers described by various experimental approach, and each set of cluster-specific marker genes were cross- referenced against published scRNAseq studies (Figure 6C and 6E) [2–4]. The cell populations identified correlated to major cell-types present in *in vivo* cardiogenesis and *in vitro* cardiac differentiation, including cardiomyocyte progenitors, cardiomyocytes, fibroblast progenitors, and cardiac fibroblasts. The TTN+ cardiomyocyte clusters, clusters 8, 9, 10, and 11 express high levels of sarcomere genes *MYH6*, *MYL3*, *TNNT2*, *TTN*. While we observe three NKX2-5 positive cardiomyocyte progenitor populations, cluster 5, 6, and 7, only cluster 5 actively expressed cell proliferation marker *MKI67*. Interestingly first heart field marker *TBX5* and second heart field marker *MEF2C* shared similar expression patterns among all cardiomyocyte progenitor clusters, suggesting these are still early-stage progenitors. We identified three Decorin (*DCN*)-positive fibroblast clusters cardiac fibroblast populations, clusters 1, 2, and 3 that contain cells expressing high levels of extracellular matrix genes such as *DCN*, *COL3A1* and *COL1A1*. We further classified cluster 1 (DCN+MKI67+ Proliferating Fibroblast) as proliferating fibroblast progenitors based on their expression of *MKi67*. Lastly, we detected an ID1+ non-cardiomyocyte progenitor cluster. In the developing heart, *ID1* is exclusively expressed by non-cardiomyocyte cells in the epicardium and endocardium [34].

Comparable cell-type clusters were detected in control and *ASXL3^fs/^*^fs^ samples when clustered independently and in aggregate. In the combined analysis, *ASXL3^fs/fs^* cells are distributed across all clusters (Figure 6C). The reproducibility of scRNA-seq transcriptomics allows the cellular composition of heterogeneous differentiating tissue to be determined between control and *ASXL3^fs/fs^* samples (Figure 6F). The ratio of non-cardiomyocytes fated cell progeny (*+/+* 20.3%, *fs/fs* 25.5%) relative to cardiomyocytes fated lineages (*+/+* 79.7%, *fs/fs* 74.5%) was similar in samples from both genotypes (Figure 6F). A slight increase in *ASXL3^fs/fs^* cardiac fibroblasts, differentiating along a *DCN* defined lineage, was observed relative to control sample (+/+ 17.8%, fs/fs 24.9%). Additional days of cardiac differentiation are required to determine if this composition difference resolves or results in a *ASXL3^fs/fs^* cardiac differentiation phenotype.

To identify transcriptomic changes that occur during cardiac differentiation, *ASXL3^fs/fs^* DEG were determined for each cluster (supplemental data). An average of 367 DEGs were detected per cluster (Min. 184; Max. 749). KEGG pathway analysis of DEGs showed overlapping dysregulation of ECM receptor interaction, signaling pathways, cardiomyopathy, and cell function related pathways (Figure 6G). These aberrant pathways were enriched for both up- and down- regulated genes across multiple clusters. Notably, three of the top KEGG pathways focused on ECM receptor interaction were showed DEG enrichment, specifically “Focal Adhesion”, “ECM- receptor interaction” and “PI3K-AKT signaling pathway” (Figure 4C and 6G). These findings suggest pathologic mechanisms are shared between mice and humans. In addition, DEGs from three NKX2-5 positive, cardiomyogenic progenitor clusters (NKX2-5+MKI67+ Progenitor, NKX2- 5+MKI67- Progenitor 1, and NKX2-5+MKI67- Progenitor 2) were similarly enriched in the Focal adhesion KEGG pathway. Of the 184, 235, and 467 DEGs identified in each cluster respectively while *MAP2K1*, *ACTB*, and *MYLK* were differentially expressed among all 3 clusters from the Focal adhesion KEGG pathway. Interestingly the majority of these DEGs were down-regulated DEGs (168, 228, 259, respectively). ECM deposited by the stromal fibroblast populations acts non-autonomously to influence cardiomyogenic progenitor proliferation (Figure 7H). Coalescence of cardiomyogenic progenitor DEGs in the focal adhesion KEGG pathway implicates that imbalanced expression of ECM ligand components and/or receptors in cardiomyogenic progenitor cells impacts proliferation and differentiation.

**Figure 7.**
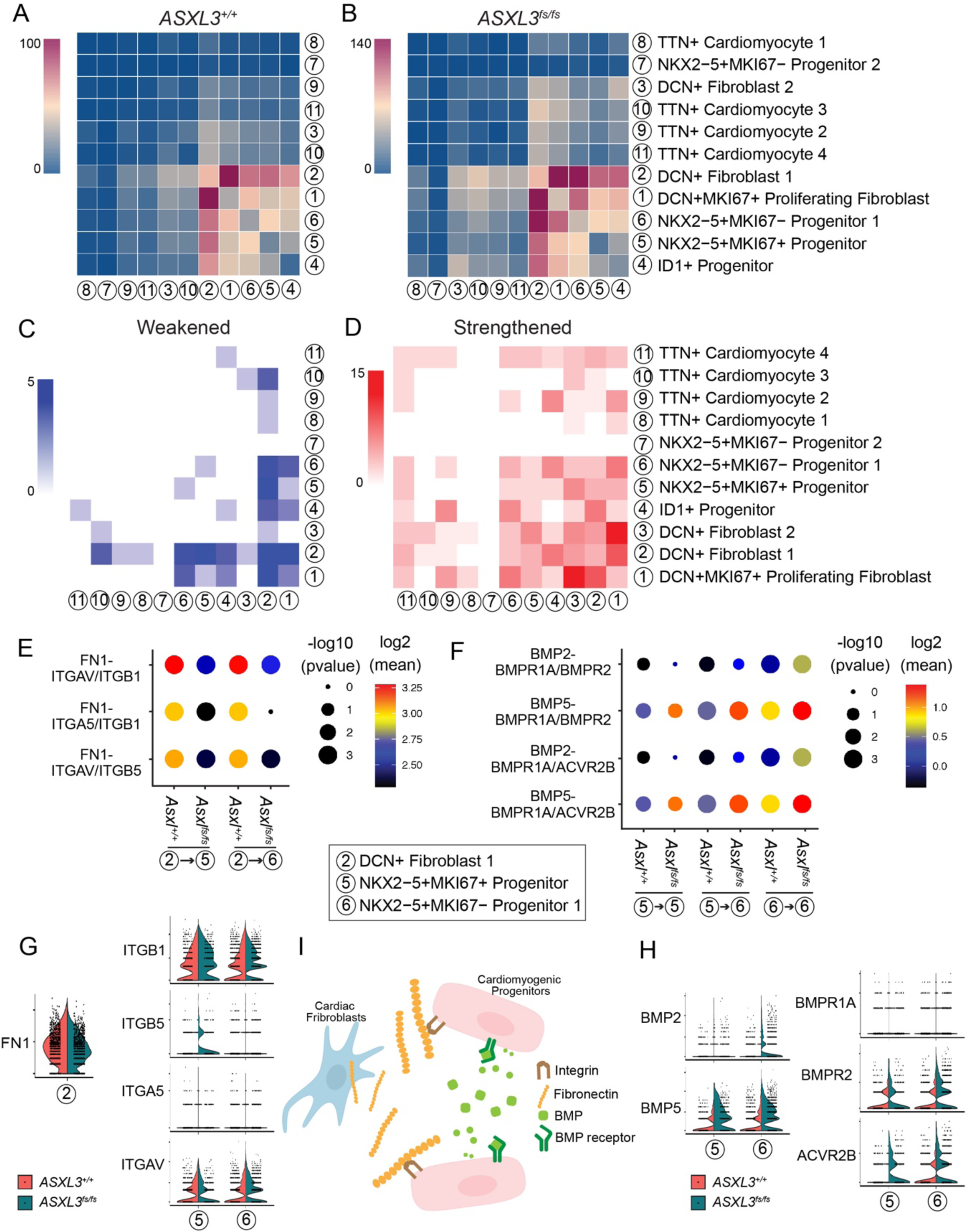
Loss of ASXL3 leads to disruptions in ECM and cell-cell communication in hESC in vitro cardiac differentiation. Heatmap of significant cell-cell signaling pathways between identified clusters in *ASXL3^+/+^* cells **A**, and *ASXL3^fs/fs^ c*ells **B**, predicted by CellPhoneDB. Note that the active communications occurred between the fibroblast clusters and the progenitor populations. Heatmap of weakened **C**, and strengthened **D**, cell-cell communication lines in *ASXL3^fs/fs^* cells compared to *ASXL3^+/+^* cells. **E,** Dotplots of Fibronectin-Integrin pathways intensities between DCN+ Fibroblast 1 and the NKX2- 5+ cardiomyogenic progenitors. Fibronectin (FN1) was expressed by DCN+ Fibroblast 1 and the NKX2-5+ cardiomyogenic progenitors expressed the Integrin receptors. CellphoneDB predicted a decrease in the *ASXL3^fs/fs^* cells. **F,** Dotplots of BMP pathways intensities between cardiomyogenic progenitor cells. Both BMP ligands and BMP receptors were expressed in cardiomyogenic progenitor populations. CellphoneDB predicted an increase in the *ASXL3^fs/fs^* cells. Gene expression violin plots of **G,** fibronectin- integrin and **I,** BMP signaling genes FN1, ITGA5, ITGAV, ITGB1, ITGB5, BMP2, BMP5, BMPR1A, BMPR2, and ACVR2B in clusters discussed in **E** and **F**. **H**, Our proposed model. From our scRNA-Seq data CellphoneDB predicted that cardiac fibroblasts expressed fibronectin, which is an important ECM substrate that supports the cardiomyogenic progenitors while the autocrine and paracrine BMP signaling occurred between the cardiomyogenic progenitors. Defects in ASXL3 led to disruptions of these cell-cell communications.

To assess defects in cell-cell communication, due to a cell-specific imbalance of ligand- receptor expression, we processed our scRNA-Seq dataset using CellphoneDB, an analysis tool developed to objectively quantify potential cell signaling between clusters based on single-cell transcriptomics [35]. The averaged expression intensity of each ligand-receptor pair was calculated and compared to the random permuted null distribution to calculate *P* values, on which are based unbiased identification of cell-cell interactions and active signaling pathways. 129 and 217 active molecular signaling pathways were identified in *ASXL3^+/+^* and *ASXL3^fs/fs^* cells respectively (Figure 7A and 7B; Figure 8 and 9 in Data Supplement) by the CellphoneDB algorithm. Interestingly regardless of genotype, DCN+ fibroblast 1 was the most “communicative” population of cells, interacting within itself and robustly with both cardiomyogenic progenitors (NKX2-5+MKI67+ Progenitor, and NKX2-5+MKI67- Progenitor 1 clusters), and non- cardiomyogenic progenitors (DCN+MKI67+ Proliferating and Fibroblast, ID1+ Progenitor) (Figure 7A and 7B). This piece of evidence again strongly favors the hypothesis that the fibroblasts support and regulate progenitors in the developing heart through both laying down the ECM environment and secreting other signaling cue molecules. Conversely in comparison mature cardiomyocytes are “quieter” and less communicative in general (Figure 7A), although in the *ASXL3^fs/fs^* condition the mature TTN+ cardiomyocyte populations were more communicative compared to their *ASXL3^+/+^* counterparts with the progenitor populations (Figure 7B). However, we also identified a group of cardiomyogenic progenitors (NKX2-5+MKI67- Progenitor 2) that had very little communication with other cells (Figure 7A). The importance and function of this group of cells remain to be further investigated.

We next compared the changes in cell-cell signaling pathways between the *ASXL3^+/+^* and *ASXL3^fs/fs^* cells. Spreadsheets of active cell-cell communication lines between each cell populations in *ASXL3^+/+^* and *ASXL3^fs/fs^* cells were generated and then cross-referenced. While changes in total of 183 different cell-cell signaling communication lines were detected, 143 were strengthened and only 40 communication lines were weakened in *ASXL3^fs/fs^* compared to *ASXL3^+/+^* control cells (Figure 7D and 7E). Strikingly, among the 40 weakened communication lines, DCN+ Fibroblast 1 was involved in 28 among the 40, suggesting an important role this group of fibroblasts play in the biology of *ASXL3^fs/fs^* defects (Figure 7C). Fibronectin 1 (FN1) was one of the most important ECM proteins downregulated by the ASXL3 loss of function identified in our mouse model. Similarly in our human in vitro model, FN1 was highly expressed in the DCN+ Fibroblast 1 cells while the different heterodimer combinations between Integrin A5/AV (ITGA5/ITGAV) and Integrin B1/B5 (ITGB1/ITGB5) were expressed by the cardiomyogenic progenitor clusters NKX2-5+MKI67+ Progenitor and NKX2-5MKI67- Progenitor 1 (Figure 10 in Data Supplement). FN1 expression was downregulated among all fibroblast clusters (Figure 10 in Data Supplement) while the integrin receptor genes were mostly downregulated in the NKX2- 5+MKI67+ Progenitor and NKX2-5MKI67- Progenitor 1 (Figure 10 in Data Supplement). CellphoneDB hence predicted a weakening of the Fibronectin (DCN+ Fibroblast 1) - Integrin (NKX2-5+MKI67+ Progenitor/NKX2-5MKI67- Progenitor 1) signaling in our *ASXL3^fs/fs^* model compared to the *ASXL3^+/+^* controls (Figure 7F and 7H). Conversely, BMP signaling was among the signaling pathways strengthened in the *ASXL3^fs/fs^* cells. BMP signaling plays a crucial role and regulates cardiogenesis during heart development and loss of one of the BMP receptor genes BMPR1A leads to decrease in cell proliferation in the developing heart [36]. Interestingly we observed that multiple BMP signaling communication lines were strengthened between and within the cardiomyogenic progenitor clusters (Fig 7F and 7I). Although we did not observe an increase in proliferating cardiomyogenic progenitors in our in vitro model, increased BMP signaling pathways may implicate proliferation defects as observed in the *Asxl3^fs/fs^* animal model (Fig 3).

## DISCUSSION

In this study, we demonstrate that pathogenic frameshift variants in *Asxl3* cause increased levels of H2AUb1, neonatal lethality, and an array of partially penetrant congenital heart defects (CHD). Hypoplasia of the right ventricle, the most prevalent *Asxl3*-associated CHD, was observed together with cellular composition changes within the heart. While different genes are dysregulated in mouse and human *ASXL3* models, both showed suppression of ECM gene sets indicating disruption of a conserved regulatory mechanism. Corresponding disruption to key signaling pathways and proliferation were detected in mouse ventricular development and human cardiac differentiation. Our findings provide an evidence for how chromatin biology can influence heart development through ECM regulation, and more broadly, a mechanism for co-occurrence of CHDs with ASD due to pathogenic variants in chromatin genes.

*ASXL1, ASXL2* and *ASXL3* are the genetic basis of Bohring-Optiz (BOS), Sashsi-Pena (SPS) and Bainbridge-Ropers syndromes (BRS) respectively [28, 37, 38], developmental disorders with multiorgan involvement. Despite a shared H2AUb1 deubiquitination molecular mechanism, each ASXL1-, ASXL2- and ASXL3-PR-DUB complex conveys non-redundant functions based on the unique constellation of clinical features attributed to each disorder. CHDs have been described in association with *de novo* monoallelic pathogenic variants in *ASXL1* (BOS) and *ASXL2* (SPS), but not *de novo* monoallelic frameshift *ASXL3* (BRS/ASD) variants. CHDs observed in individuals with BOS include ventricular and atrial septal defects with minor incidence of cardiac hypertrophy and bradycardia. SPS is characterized by atrial septal defects, patent ductus arteriosus (PDA) and left ventricular dysfunction. While the pathogenicity of monoallelic *ASXL3* missense variants is difficult to classify, CHDs were recently reported for biallelic *ASXL3* missense variants, providing evidence of the pathogenicity of *ASXL3* missense variants [30]. Distinct CHDs detected in loss-of-function (knockout or frameshift) *Asxl1*, *Asxl2* and *Asxl3* mouse models, support non-redundant roles for ASXL family members in heart development, while highlighting the discrepancy in dosage-sensitivity between the species [27, 39].

Cardiac phenotypes of biallelic loss-of-function *Asxl* mouse models mirror the diversity of clinical features observed across the ASXL-related developmental disorders. Highly penetrant severe inlet ventricular septal defects, with interventricular septum (IVS) involvement are observed in *Asxl1* knockout mouse model, reminiscent of cardiovascular features of BOS [27]. Minor membranous VSDs were detected in *Asxl2*, while muscular VSDs were found in *Asxl1* and *Asxl3* mice. The severe *Asxl1* VSD phenotype can also be distinguished from *Asxl2^-/-^* and *Axsl3^fs/fs^* heart phenotypes by normal ventricular wall and valve thickness. Partially penetrant ventricular hypoplasia is observed in both *Asxl2^-/-^* and *Asxl3^fs/fs^* mouse models, but exhibit distinct ventricular laterality involvement. *Asxl2^-/-^* animals are characterized by hypoplastic left ventricle and IVS with corresponding thickening of the compact myocardium, while *Asxl3^fs/fs^* animals present with right ventricular hypoplasia. The general contribution of FHF and SHF cardiac primordia, to the left versus right ventricles respectively, highlight the spatiotemporal specificity of the individual *Asxl2* and *Asxl3* family members. Identifying the non-redundant genome-wide H2A deubiquitination functions of individual PR-DUB complexes will be critical future studies to correlate direct genomic targets of individual PR-DUB complexes to mechanisms of heart development.

Based on the cardiac involvement in the *Asxl3^fs/fs^* mouse, we analyzed echocardiograms of three individuals clinically diagnosed with BRS based on *de novo* frameshift *ASXL3* variants, which revealed a single case of bicuspid aortic valve. Furthermore, review of published case reports revealed one additional BRS case displaying cardiac phenotypes as pulmonary artery stenosis, small patent foramen ovale and PDA [38, 40]. This analysis did not investigate digenic or co-occurrence of a second pathogenic coding variants in these cases, yet the incidence of these CHDs is insufficient evidence to conclude a direct link between *de novo* monoallelic frameshift *ASXL3* variants and CHDs. Individuals with biallelic missense *ASXL3* variants present with Tetralogy of Fallot (TOF) [30]. TOF refers to the combination of four congenital abnormalities; VSD, pulmonary valve stenosis, misplaced aorta and thickened right ventricular wall. Thickened right ventricular wall and VSD phenotypes phenocopy the CHDs detected in *Asxl3fs/fs* mouse alluding to a consistent genotype-phenotype correlation. The developmental mechanism implicated by transcriptomic analysis of a biallelic missense *Asxl3* mouse model implicated involvement of the PI3K/AKT KEGG pathway, also detected in our *Asxl3^fs/fs^* model, and altered expression of components of the PRC2 complex, a phenotype shared with the *Asxl2^-/-^* mouse model [27, 30, 39]. These phenotypic outcomes provide further evidence for the importance of chromatin biology and Polycomb transcriptional repression in heart development.

The occurrence of heart defects in mouse models that disrupt genes encoding PRC1, PR- DUB, and PRC2 components demonstrates the importance of Polycomb transcriptional plasticity during cardiogenesis [22]. Phenotypic overlap between PR-DUB and PRC1 models, as compared to PRC2, implicates dynamic exchange of distinct Polycomb histone PTMs as central to the cardiac morphogenesis phenotypes [27, 41–43]. Similar to the *Asxl3^fs/fs^* mouse model, right ventricular hypoplasia was detected in the knockout mouse model of *Phc1* (also known as *Rae28)*, which encodes canonical PRC1 complex (PRC1.2 and PRC1.4) component PHC1 [42]. VSDs, reminiscent of the cardiac phenotype observed in the *Asxl1^-/-^* mouse model, have been described in *Phc1* and the noncanonical PRC1 (PRC1.1) complex component *Bcor* knockout mouse models [44, 45]. While dynamic exchange of H2A mono-ubiquitination by PRC1 and PR- DUB complexes are clearly essential for cardiac development, careful evaluation of cardiac morphogenesis for defects associated with a comprehensive representation of PRC1 complexes is required to assign non-redundant roles in heart development and complementary PR-DUB phenotypes. Alternatively, PRC2 heart phenotypes are distinct from those shared between mouse models of PRC1 and PR-DUB components, highlighting the paucity of data detailing how H2AUb1 and H3K27Me3 dependent transcriptional plasticity is coordinated. Knockout mouse models of PRC2 components *Ezh2* or *Jarid2* present with a spectrum of cardiac structural malformations including hyper-trabeculation and hypoplasia of the compact myocardium, which is characterized by thinning of the ventricular wall [21, 46–48]. Neither do PRC2 associated heart phenotypes exhibit phenotypic laterality, by preferentially disrupting morphogenesis of a single ventricle.

Genetic and functional studies have contributed to our current understanding of the pathogenic mechanisms that give rise to single ventricular disorders (SVDs), namely hypoplastic left heart syndrome (HLHS) and hypoplastic right heart syndrome (HRHS). HRHS is a phenotype shared with the *Asxl3^fs/fs^* mouse model. Both HLHS and HRHS are characterized by underdeveloped or malformed structures of a single ventricle. Human and mouse genetic studies of SVDs have identified chromatin regulators, transcription factors, and signaling pathway components as the basis of these developmental defects [42, 49–62]. The convergence of the human genetics and animal models on these common molecular pathways implicate a shared pathogenic mechanism that impinges on CPC proliferation and ECM composition [52-54, 63, 64].

The left and right ventricle have distinct FHF versus SHF origins. Nevertheless, mouse HLHS- relative to HRHS ventricular transcriptomics not only reveal DEGs enriched in similar signaling pathways, but implicated altered ECM based mechanisms. The enriched KEGG pathways include ECM receptor interaction, Calcium signaling pathway, Focal adhesion, TGF-Beta signaling, Dilated cardiomyopathy, Hypertrophic cardiomyopathy, and Arrhythmogenic RV cardiomyopathy [54]. Consistent with these findings, analysis of ECM content in left and right ventricles of HLHS post-mortem and myocardial biopsy samples revealed reduced collagen levels compared to age- matched controls [64, 65]. Together these results suggest altered ECM dynamics contribute to laterality of ventricular hypoplastic heart phenotypes, but does not provide a mechanism for how asymmetric expressivity is established.

Genes that encode ECM components are differentially expressed in both ASXL3 mouse and human models. Developmentally, the ECM functions as a substrate for cell migration and promotes CPC proliferation, differentiation, and maturation, chamber formation, and valve development [66]. ECM composition is tailored, spatially and temporally to promote chamber- specific signaling pathways and structural integrity to direct these morphogenetic events. ECM coordination of cell-cell communication in cardiac development is well established, and exemplified by the influence of FN1, expressed by cardiac fibroblasts, on proliferation of CPCs through *β*1 integrin receptor binding to FN1 and PI3K/Akt pathway signaling. The importance of dynamic remodeling of ECM composition for organogenesis has been acknowledged, where disrupting either synthesis or degradation of the ECM results in CHDs attributed to disorganization of the dynamic tissue morphogenesis [15, 16, 66, 67]. More recent studies have described the network of transcription factors and chromatin-based regulatory mechanisms complexes that spatially and temporally coordinate expression of ECM genes [21, 68, 69]. For instance, conditional endocardial specific knockout of *Brg1*, a chromatin remodeler, allows for increased expression of the matrix metalloproteinase ADAMTS1, which results in premature degradation of the cardiac jelly and excessive trabeculation [12]. Our data provides additional evidence to link chromatin biology to the regulation of ECM components during cardiogenesis.

Expression of *Asxl3* at single-cell resolution during cardiac development has not been determined. *Asxl3* expression has been documented, from single-cell transcriptomics, in intermediate cell types along the Nkx2-5 cardiomyocyte lineage, and in cardiac fibroblasts of fetal mouse heart [5, 70]. Cardiac fibroblasts are an important source of diverse secreted components of the ECM [71]. In the *Asxl3* model, we detect fewer vimentin-positive fibroblasts in P0.5 *Asxl3^+/fs^* and *Asxl3^fs/fs^* ventricles and reduced expression of genes that encode ECM. We hypothesize that dysregulation of ECM during heart development contributes to E13 CPC reduced proliferation, that across development preferentially impinges on right ventricular development. Our analysis does not allow us to predict the initial source of this molecular pathology; we cannot preclude that loss of Asxl3 alters the differentiation of cardiac lineages that disrupts ECM composition and/or Asxl3-dependent ECM composition changes alter production of CPCs during development. We discovered extensive overlap between KEGG pathways enriched in our mouse model and other HLHS mouse models that centered around ECM composition, signaling, and cardiac defects [54]. The pathways included ECM receptor interaction, calcium signaling pathway, focal adhesion, TGF-Beta signaling, dilated cardiomyopathy, Hypertrophic cardiomyopathy, and Arrhythmogenic RV cardiomyopathy. Similar disrupted pathways are also observed in the biallelic *Asxl3* missense mouse [30]. Two models for SVDs have been proposed to explain this phenotype. One attributes decreased blood flow during a critical period of ventricular development to the reduced ventricular size [72]. The second model suggests CPC hyperplasia as the driving force for underdeveloped ventricles [51]. The increased proliferation we observe in *Asxl3^fs/fs^* ventricles provides further evidence for the second model, but cannot rule out a role for hemodynamics as it was not assessed.

Interestingly, molecular mechanisms implicated from the *Asxl3* mouse model were generally corroborated by *in vitro* human cardiac differentiation of bi-allelic *ASXL3* frameshift hESC lines. In both species, phenotypic differences were attributed to changes in ratio of CPC subtypes and ECM mediated signaling defects, while the distinct genetic and molecular details of each pathogenic mechanism are species-specific. In the human *in vitro* model system, the ratio of cardiac fibroblast lineages, relative to NKX2-5 cardiomyocyte fated lineages were greater in *ASXL3^fs/fs^* cultures relative to controls. This is opposite to the trend observed in the *Asxl3* mouse model where a reduction in cardiac fibroblasts were observed. The significance of the opposing observation is difficult to conclude due to differences between species, in the stage of differentiation/development, and *in vitro* culture relative to *in vivo* development. Conversely, ECM components, such as members of the collagen super family and FN1 were prominently differentially expressed in both models. While assessment of cell-cell interaction, using single cell transcriptomics, specifically indicated weakened cardiac fibroblast FN1 signaling interaction via cardiomyocyte progenitor expressed ɑv*β*1, ɑ5*β*1, and ɑv*β*5 integrin, a reduction in FN1 expression was also detected in bulk RNA sequencing results from the mouse model (Figure 4A and 7I).

Reduced FN1 expression has also been reported upon loss of HLHS associated genes ETS1, CHD7, and KMT2D in endocardial cells and may represent shared pathology among chromatin modifying genes [20].The influence of FN1 on the mitotic behavior of cardiomyocytes depends upon FN1/*β*1 integrin signaling through the PI3K/Akt pathway, a features shared by all ASXL3 models to date [14, 20].Collagen subtypes also fulfill structural and signaling cohesiveness to ECM functions and fall into broad functional groups. Four collagen subfamilies were downregulated in the mouse (1,4,5 &6), while substantially more subfamilies were differentially expressed in the human model (1,4,5,6,8,9,11,12, 13, 14, 18 & 25) (Fig 4A,B and 6G). This evidence suggests that ASXL3 has a conserved role in regulating ECM cardiac biology and highlights the importance of this cell biology for normal and pathogenic cardiogenesis. Nevertheless, it will be important to resolve if altered differentiation of cardiac lineages initiate a cascade of ECM composition changes and cell-cell signaling defects, or if chromatin changes directly alter expression of ECM components.

Human genetic studies have described the high incidence of pathogenic de novo variants in chromatin genes that are the genetic basis of syndromic ASD with partially penetrant CHDs [50, 73, 74]. *ASXL3* is a high confidence ASD gene [75]. *De novo* monoallelic frameshift *ASXL3* alleles, also the genetic basis of BRS, are also recognized as the genetic basis of ASD. The low to high penetrance of CHDs between *Asxl3^+/fs^* to *Asxl3^fs/fs^* genotypes harkens the co-occurrence of ASD and CHDs and suggests that reduced penetrance of CHDs in ASD may reflect increased vulnerability of developing brain relative to heart to de novo pathogenic variants in chromatin genes. Future studies will be required to determine if this is due to organ-specific differences in dosage sensitivity of chromatin genes and/or represents differences in organ-specific thresholds at which altered histone modifications genome-wide impact organ development.

## METHODS

Detailed methods regarding animal model and human cardiomyocytes are described in the Data Supplement. All sequencing data sets in this article are deposited in international public repository, Gene Expression Omnibus (GEO), under accession IDs for mouse and human cardiomyocytes bulk RNA sequencing and GSE for single cell RNA Sequencing from human cardiomyocyte.

### Animals

All experiments were performed in accordance with animal protocols approved by the Unit for Laboratory Animal Medicine (ULAM), University of Michigan. A detailed description for the generation of the *Asxl3^fs/fs^* mouse line is in the Data Supplement. Briefly, CRISPR/cas9 was used to edit a region in *Asxl3* exon 12. The differential *Asxl3* expression was validated by qRT-PCR and western blot. *Asxl3^+/fs^* mice were maintained on a C57BL/6 background. Heterozygous breeding was used for experiments with E0.5 established as the day of vaginal plug.

### Histology and morphometric measurements

Mouse tissues were fixed for 24h in 4% paraformaldehyde. Whole-mount skeletal preparations were stained with Alician blue and Alizarin red (Data Supplement). Whole bodies were processed for paraffin embedding, sectioned at 5-µm intervals and stained with Hematoxylin/eosin or Masson’s trichrome. ImageJ was used to measure interventricular septum (IVS), left ventricle lumen width (LVLW), left wall thickness (LWT), right ventricular lumen width (RVLW), and right wall thickness (RWT) from 9 *Asxl3^+/+^*, 4 *Asxl3^+/fs^* and 14 *Asxl3^fs/fs^* P0.5 hearts.

### Immunohistochemistry

Detailed descriptions for our immunohistochemical studies are in the Data Supplement. 13-um thick cryosections were incubated with the listed primary and secondary antibodies. EdU- positive cells were detected in cryosections with the Click-IT EdU imaging kit (Invitrogen, Carlsbad, CA).

### RNA sequencing

Total RNA was isolated from ventricles of 4 *Asxl3^+/+^*, 4 *Asxl3^+/fs^*, and 4 *Asxl3^fs/fs^* littermates. Single-end library was prepared as described in the Data Supplement and sequenced with the Illumina HiSeq 2500. Single-end sequenced reads were aligned to the mouse reference genome (mm10) using Tophat2 (Tophat 2.0.4) with default settings. Fragment quantification was computed using feature Counts and annotated according to RefSeq genes. DESeq2 was used to calculate estimates of dispersion and logarithmic fold changes to perform the expression normalization and differential expression analysis. DAVID analysis was performed with up- and down-regulated genes to identify enriched KEGG pathways (Huang et al. 2009). Gene set enrichment analysis was performed using the C5 ontology gene set from the molecular signature database [76].

### Generation and differentiation of *ASXL3fs/fs* hESCs

To generate the patient variants identified by our lab [29](Srivastava et al. 2016) in hESCs, we designed sgRNAs to a region flanking the *ASXL3* variant (c.1443dupT). We modified a previously described small-molecule WNT inhibitor protocol to differentiate hESCs toward a cardiac lineage (Data Supplement).

### SEQ-well single-cell RNA sequencing and analysis

Single cell suspensions of *ASXL3^+/+^* (n=3) and *ASXL3^fs/fs^* (n=2) hESCs were obtained after 12 days of cardiac differentiation. Isolated cells were loaded onto a SEQ-well array and processed as described (Data Supplement). Single-cell libraries were sequenced on an Illumina NextSeq 75 cycle instrument. Reads were processed following the Drop-seq core computational protocol and quality controlled. We integrated all the *ASXL3^+/+^* and *ASXL3fs/fs* samples with the “Find Integration Anchors” and “Integrate Data” functions of Seurat (v3.1.2). Using Seurat, we performed Uniform Manifold Approximation and Projection (UMAP) dimensionality reduction with the integrated data to identify cell-type clusters. We performed Wilcoxon-rank sum tests to identify differentially expressed genes between genotypes in each cluster. DAVID analysis was performed with up- and down-regulated genes to identify enriched KEGG pathways. CellPhoneDB was used to predict cell-to-cell communications [35].

### Statistics

We analyzed data using GraphPad Prism with values being represented as means±SEM. Student’s 2-tailed *t* test was used to generate *P-* values. Chi square analysis was used for statistical analysis of Mendelian ratios. *P-* values <0.05 were considered significant. Benjamini- Hochberg multiple hypothesis corrections of the *p* values was used to test significance after DAVID analysis. Wilcoxon rank-sum test was used to identify differentially expressed genes from our scRNA-seq.

## SUPPLEMENTAL METHODS

### Single-guide RNA (sgRNA) design

Potential sgRNAs for the target sequence in *Asxl3* gene were identified by a previously published scoring algorithm (http://crispr.mit.edu). Selection was made on the basis of a qualitative balance of specificity scores, distance to desired mutation/insertion and manual assessment of the off-target list. A total of three sgRNAs were selected. Bicistronic expression vector px330 expressing Cas9 and sgRNA [77] was used for cloning the sgRNAs (Addgene) as described previously [78].

### Genotyping

A product size of ∼850 bp flanking the *Asxl3* mutation is generated by PCR using primer pair, forward: TCACATGGCTTAGTGGTTGT, reverse: CTGTTCTTCGGGGTCACTCT. Sanger sequencing and ligation detection reaction are used for genotyping of CRISPR/Cas9 edited *Asxl3* alleles [79].

### Off-target mutation analysis

In the off-target list assessment, we considered the preference to avoid sgRNAs with potential hits in coding regions, sgRNAs with off-target hits on the same chromosome as the intended target, and, when possible, any sgRNAs that had many predicted off-targets lacking mismatches in the seed region (10–12 nucleotide proximal to the protospacer-adjacent motif (PAM), as well as whether off-targets had NGG or NAG PAMs. Predicted off-target sites included regions in *Sdc* and *Gsta1.* Two approaches were used to validate the on-target and off-target CRISPR/Cas9 genome-editing. One, Sanger sequencing of the predicted on-target and off-target regions and two, whole genome sequencing (WGS) were performed. No predicted pathogenic single nucleotide polymorphisms, indels or structural variants such as inversions, rearrangements, duplications and major deletions were detected (Figure 1 in Data Supplement).

### Pronuclear injection

Sexually immature female C57BL/6 mice (4 weeks old) were superovulated by intraperitoneal injection of 5 IU Pregnant mare serum gonadotropin (PMSG) followed by 5 IU human chorionic gonadotropin (hCG) at an interval of 48 h and mated overnight with C57BL/6 male mice that were >12 weeks old. Fertilized eggs were collected after 20 hours of hCG injection by oviductal flashing, and pronuclei-formed zygotes were put into the M2 medium. Microinjection was performed using a microinjector (Narishige) equipped microscope. The injected eggs were cultivated overnight in potassium simplex optimization medium (KSOM) at 37C 5% CO2 humidified incubator. Two-cell stage embryos were transferred into the oviducts of 0.5 days pseudopregnant C57BL/6 X DBA/2 F1 females. After birth, ∼1 mm tail biopsy from 2–4-day-old pups was used as a source of DNA for *Asxl3* genotyping (forward: TCACATGGCTTAGTGGTTGT, reverse: CTGTTCTTCGGGGTCACTCT).

### Western blot analysis

Complete hearts from E13 *Asxl3+/+* and *Asxl3fs/fs* mice were homogenized in RIPA buffer supplemented with protease inhibitor cocktail and phosphatase inhibitor cocktail 3 obtained from Sigma-Aldrich (P8340 and P0044; St Louis, MO, USA). Protein levels were normalized after BCA analysis (Pierce). Cell lysates were separated using electrophoresis on 4-20% SDS- polyacrylamide gels and transferred to PVDF membrane (Millipore, Billerica, MA, USA). For western blot, after the transfer, the PVDF membrane was blocked with 5% milk and incubated with following antibodies overnight. Primary antibodies used: anti-ASXL3 (Bielas Lab, 1 to 200), anti-ubiquityl-Histone H2A (Cell Signaling Technology, 8240, 1 to 2000), anti-Histone H3 (Abcam, Ab10543, 1 to 5000). Donkey anti-rabbit HRP-conjugated (Cytiva, NA9340V, 1 to 5000) and goat anti-mouse HRP-conjugated (Invitrogen, 32430, 1 to 10000) were used for 1h incubation at room temperature. Antibody incubation and chemiluminescence detection were performed according to manufacturer’s instruction [ThermoFisher Scientific (Waltham, MA, USA) cat no. 34095].

### Skeletal preparations

For skeletal preparations, newborn animals were skinned and fixed in 95% ethanol. Fixed skeletons were stained with Alcian Blue (76% ethanol:20% acetic acid) at 37°C for 48 h, rinsed in 95% ethanol, treated with 1% KOH for 4-5 h and stained with Alizarin Red in 2% KOH for 1 h. Stained skeletons were cleared successively in 20% glycerol:1% KOH, 50% glycerol:1% KOH and 100% glycerol (n = 7 for each genotype). Forelimbs were removed and imaged on a Leica MZ125 stereomicroscope.

### H&E and Masson’s Trichrome staining

Newborn mice were sacrificed and the whole body was fixed for 24 h in 4% paraformaldehyde, dehydrated, and embedded in paraffin. Paraffin blocks were serially sectioned at 5 µm thickness and stained with Hematoxylin/eosin and Masson’s trichrome.

### Immunohistochemistry

Hearts were dissected and removed from mice at E14 or P0.5 and then kept in 4% PFA at 4℃ overnight. Hearts were cryopreserved by submersion in 20% then 30% sucrose solutions and embedded in OCT cryosectioning media (Tissue-Tek, Torrance, CA). 13 μm cryosections were obtained. After thawing, sections were incubated with PBS for 15 min to wash away OCT. For antibodies that required antigen retrieval, cryosections were heated in 10 mM sodium citrate for 20 minutes at 95℃ followed by incubation at room temperature for 20 minutes and 3 PBS washes. Sections were then incubated with a normal donkey serum (NDS) blocking buffer [5% NDS (Jackson ImmunoResearch), 0.1% Triton X-100, 5% BSA] for 1 hour. Subsequently, they were incubated with primary antibodies diluted in NDS blocking buffer at 4℃ overnight, washed with PBS, and stained with secondary antibodies at room temperature for 1 hour. Slides were washed with PBS, incubated with DAPI for 5 minutes and coverslipped with MOWIOL. The following antibodies and dilutions were used: COL1A1 (Novus Biologicals, NB600-408, 1 to 500), COL3A1 (Abcam, ab7778, 1 to 100), VIM (Abcam, ab195878, 1 to 500). AlexaFluor-conjugated secondaries were: donkey anti-rabbit 647 (Invitrogen, A31573, 1 to 400), anti-rabbit 555 (Invitrogen, A31572, 1 to 400), donkey anti-mouse 555 (Invitrogen, A31570, 1 to 400), donkey anti-mouse 647 (Invitrogen, A31571, 1 to 400).

### Image analysis and quantification

Images of immunostained slides were acquired with a Nikon A1 confocal microscope and processed with LAS X software. Positively stained cells from anatomically-matched serial sections were quantified with ImageJ software. Scaled images were segmented using the grid function in ImageJ with the area per point set to 50000 pixels^2 for each image. Positively stained cells within a segment were counted using the ImageJ Cell Counter plugin. To measure the cross- sectional area of cardiomyocytes we used wheat germ agglutinin Alexa Fluor™ 488 Conjugate (Abcam, W11261, 1 to 100). ImageJ software was used to calculate the cross-sectional area of the cardiomyocyte.

### EdU birth-dating analysis

Females from timed-pregnant matings were injected with EdU (20 mg/kg) at embryonic day 13. 24 hours later, hearts were dissected and removed from EdU injected embryos then kept in 4% PFA at 4℃ overnight. EdU labeling was detected in cryosections by using the Click-IT EdU imaging kit (Invitrogen, Carlsbad, CA) according to the manufacturer’s instructions. After sections were incubated with Click-IT reaction cocktail, they were washed with normal donkey serum blocking buffer and then additional antibody staining was performed.

### E18.5 heart ventricular RNA sequencing

Total RNA from E18.5 ventricles was extracted from *ASXL3+/+*, *ASXL3+/fs*, and *ASXL3fs/fs* littermates. Transcriptome libraries were prepared using 200–1000 ng of total RNA. PolyA + RNA isolation, cDNA synthesis, end-repair, A-base addition and ligation of the Illumina indexed adapters were performed according to the TruSeq RNA protocol (Illumina). Libraries were size selected for 250–300 bp cDNA fragments on a 3% Nusieve 3:1 (Lonza) gel, recovered using QIAEX II reagents (QIAGEN) and PCR amplified using Phusion DNA polymerase (New England Biolabs). Total transcriptome libraries were prepared as above, omitting the poly A selection step and captured using Agilent SureSelect Human All Exon V4 reagents and protocols. Library quality was measured on an Agilent 2100 Bioanalyzer for product size and concentration. Single-end libraries were sequenced with the Illumina HiSeq 2500 with sequence coverage to 100-150 m reads. Single-end sequenced reads were aligned to the mouse reference genome (mm10) using Tophat2 (Tophat 2.0.4) with default settings. Fragment quantification was computed using feature Counts and annotated according to RefSeq genes. DESeq2 was used to calculate estimates of dispersion and logarithmic fold changes to perform the expression normalization and differential expression analysis. Further genes and isoforms were annotated with NCBI Entrez GeneIDs and text descriptions. We used gene ontology consortium (http://geneontology.org) and the DAVID database (https://david.ncifcrf.gov/) for the enrichment analysis of the set of DEGs to identify significantly enriched functional categories. Benjamini-Hochberg multiple hypothesis corrections of the *p* values was used, and *p* values less than 0.05 were called to be significant changes.

### hESC CRISPR/Cas9 editing

Human embryonic stem cell line H9 (WiCell) was maintained in mTeSR1 media (Stemcell Technology) on matrigel coated tissue culture dishes. Undifferentiated human Embryonic Stem Cells (hESCs) hESCs were passaged three times every two weeks with daily media change. To generate the patient equivalent variants identified by our lab [29] in hESCs, we designed the sgRNAs flanking the *ASXL3* variant (c.1443dupT). Pluripotent H9 cells were electroporated with PX330 plasmids (described above) with Amaxa electroporator using manufacturers protocol (Lonza). After electroporation single cells were plated in 96 wells and viable healthy colonies were then sequenced to confirm the gene editing in *ASXL3*.

### Cardiac differentiation of hESCs

We modified previously described small-molecule protocols to differentiate hESCs toward a cardiac lineage [32, 80]. In brief, 1x10^6 hESCs were seeded per well in matrigel coated 6-well tissue culture plates. Once the culture reached 100% confluence within 3 days the media was changed to RPMI 1640 with 1X B27 supplement minus Insulin (RPMI/B27- media) with 6 uM CHIR99021 and the day marked as differentiation day 0. 24 hours later the media was changed back RPMI/B27- media without CHIR99021 for another 24 hours. Afterwards CHIR99021 was added back for another 24 hours. On day 3 the old media was removed and washed once with PBS. The cells were then incubated in RPMI/B27- with 5 uM Wnt inhibitor IWP-4 for additional 48 hours. At the conclusion of Wnt inhibition the media was changed back to RPMI/B27- and replaced every other day until day 10. After day 10 the media was changed to RPMI 1640 with 1X regular B27 Supplement (RPMI/B27+ media) with media replacement every other day. Cardiomyocytes typically start contracting between day 7 and day 12. If no contracting cardiomyocytes were observed by day 12, the cells were discarded, otherwise the cells were dissociated and subject to single cell RNA sequencing.

### Seq-Well single-cell RNA-sequencing

Seqwell was performed as described [81, 82]. Briefly, functionalized Seq-Well arrays, containing 90,000 picowells, were loaded with barcoded beads (ChemeGenes, Wilmington, MA). 20,000 cells were loaded onto the arrays and incubated for 15 minutes. To remove residual BSA and excess cells, arrays were washed with PBS. Functionalized membranes were applied to the top of arrays, sealed in an Agilent clamp, and incubated at 37° for 45 minutes. Sealed arrays were incubated in a lysis buffer (5 M guanidine thiocynate, 1 mM EDTA, 0.5% sarkosyl, 1% BME) for 20 minutes followed by a 45 minute incubation with hybridization buffer (2 M NaCl, 1X PBS, 8% PEG8000). Beads were removed from arrays by centrifuging at 2000xg for 5 minutes in wash buffer (2 M NaCl, 3 mM MgCl2, 20 mM Tris-HCl pH 8.0, 8% PEG8000). To perform reverse transcription, beads were incubated with the Maxima Reverse Transcriptase (Thermo Scientific) for 30 minutes at room temperature followed by overnight incubation at 52℃. Reactions were treated with Exonuclease 1 (New England Biolabs) for 45 minutes at 37℃. Following we performed a second strand synthesis reaction with the Maxima Reverse Transcriptase (Thermo Scientific) for 1 hour at 37℃ as described (Hughes et al. 2020). Whole transcriptome amplification was performed using the 2X KAPA Hifi Hotstart Readymix (KAPA Biosystems). Beads were split to 1,500-2,000 per reaction and run under the following conditions 4 Cycles (98℃, 20s; 65℃, 45s; 72℃, 3m) 12 Cycles (98℃, 20s; 67℃, 20s; 72℃, 3m) final extension (72℃, 3m, 4℃, hold).

Products were purified with Ampure SPRI beads (Beckman Coulter) at a 0.6X volumetric ratio then a 1.0X volumetric ratio. Libraries were prepared using the Nextera XT kit (Illumina) and libraries were sequenced on an Illumina NextSeq 75 cycle instrument.

### Seq-Well data preprocessing

Sequencing reads were processed into a digital gene expression matrix using Drop-seq software as described [83]. FASTQ files were converted into bam files before being tagged with cell and molecular barcodes and trimmed. After converting back to FASTQs, reads were aligned to hg19 with STAR. BAM files are then sorted, merged, and tagged with gene exons. Bead synthesis errors were corrected as described and digital gene expression matrices were generated. For downstream analysis Cells with fewer than 300 detectable genes, greater than 5000 genes or greater than 10% mitochondrial genes were removed. Genes that were detected in less than 5 cells were also excluded. A total of 9,379 cells captured from *ASXL3^+/+^* and 9,198 cells captured from *ASXL3^fs/fs^* cultures passed the quality control and were used in the final analysis.

### UMAP dimensionality reduction and cluster annotation

We used the Seurat package (v3.1.2) to perform dimensionality reduction. We used the integrated and normalized data as the input to the RunPCA function of Seurat (v3.1.2) in order to compute the first 100 PCs. After that, we used the elbow algorithm to find the optimal number of PCs to construct Uniform Manifold Approximation and Projection (UMAP) plots. Visualizations in a two-dimensional space were done using RunUMAP function of Seurat (v3.1.2) for the integrated data using previous dimensional reduction data and predetermined best PC number. We performed a graph-based clustering approach using FindNeighbors and FindClusters functions of Seurat (v3.1.2). A K-nearest neighbor graph was constructed based on the euclidean distance in the predetermined PCA dimension with drawn edges between cells with similar expression levels and then refined the edge weights between any two cells. We then clustered the cells based on modularity optimization technique: Louvain algorithm with resolution parameter 0.5 and partitioned the graph constructed before into communities. We then collected cluster marker genes using the Wilcoxon rank-sum test which is a nonparametric test between the cells in a single cluster and all other cells with log fold change threshold as 0.2. To assign identities to clusters, we cross-referenced the marker genes with previously described cardiac subtype markers [2–4].

### Single-Cell differential gene expression and pathway enrichment analysis

We used the FindMarkers function in Seurat and performed a Wilcoxon rank-sum test (logFC.threshold=0.15) to compute differentially expressed genes between *ASXL3^+/+^* and *ASXL3^fs/fs^* cells within each cluster. An P-value cut-off < 0.05 was used to identify differentially expressed genes. DAVID analysis (https://david.ncifcrf.gov/tools.jsp) was performed with the up-, down-regulated, and combined genes to identify enriched KEGG pathways.

### CellPhoneDB analysis

CellPhoneDB V2 was used to predict changes between genotypes in cell-to-cell communication. *ASXL3^+/+^* and *ASXL3^fs/fs^* were analyzed individually using the metadata from the integrated Seurat object as described [35]. Ligand-receptor pair expression means and pvalues were calculated. For comparison between *ASXL3^+/+^* and *ASXL3^fs/fs^* pvalues of each ligand- receptor pair expression score in both genotypes have to be below 0.05 to be considered. Significant ligand-receptor pair signaling communication line changes in *ASXL3^fs/fs^* were identified as gained if *ASXL3^+/+^* mean=0 and *ASXL3^fs/fs^* mean >0; lost if *ASXL3^+/+^* mean>0 and *ASXL3^fs/fs^* mean =0; strengthened if (*ASXL3^fs/fs^* mean/*ASXL3+/+* mean) >1.5; or weakened if (*ASXL3^+/+^* mean/*ASXL3^fs/fs^* mean) >1.5.

## ACKNOWLEDGMENT

### SOURCES OF FUNDING

This work was supported by the Eunice Kennedy Shriver National Institute of Child Health and Human development (R01AWD010411 to SLB), National Institute of Neurological Disorders and Stroke (R01NS101597 to SLB) Simons Foundation Directors Award (19-PAF06833,17- PAF05917 SLB), Leo’s Lighthouse Foundation to SLB and BTM, NIH Cellular and Molecular Biology Training Grant ( T32-GM007315 to BTM), Genetics Training Grant (to AM). Department of Biotechnology, Innovative Young Biotechnologist Award (BT/12IIYBAl2019/13) and Ramalingaswami Fellow (<u>D.O.NO.BT/HRD/35/02/2006</u>) to AS.

## Nonstandard Abbreviations and Acronyms

ASXL3: Additional sex comb like 3
BRS: Bainbridge Ropers Syndrome
sc-RNA: seq Single-cell RNA sequencing
PR-DUB: Polycomb Repressive-Deubiquitination
CHDs: Congenital Heart Defects
CM: Cardiomyocytes
CF: Cardiac fibroblasts
sgRNA: Single-guide RNA

---

**Supplemental Figure 1.**
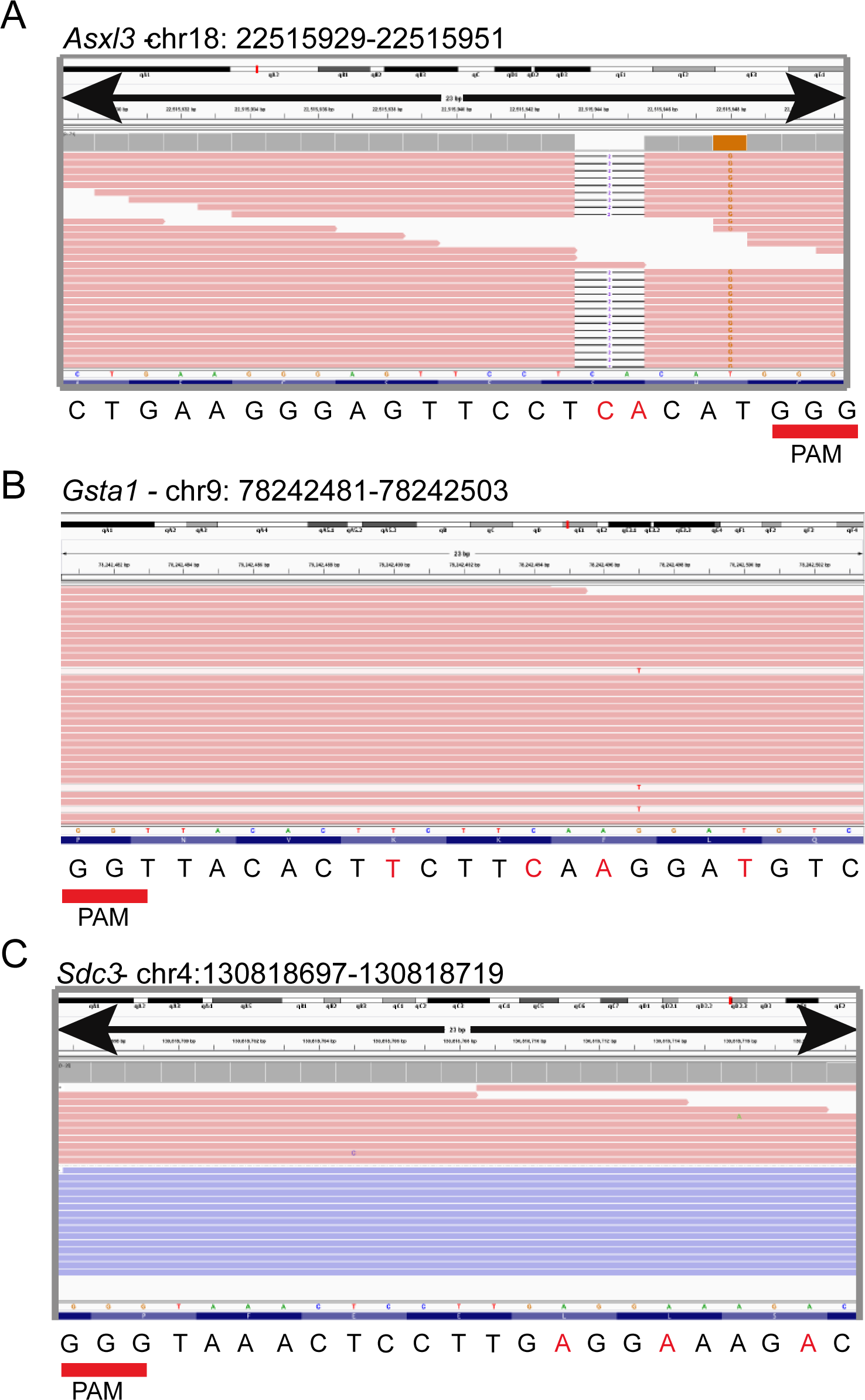
Whole Genome Sequencing to confirm CRISPR editing of Asxl3. IGV viewer windows of whole genome sequencing results from Asxl3^fs/fs^ mice. Displayed are sgRNA target site regions for **A,** *Asxl3*, **B,** *Gsta1*, and **C,** *Sdc3*.

**Supplemental Figure 2.**
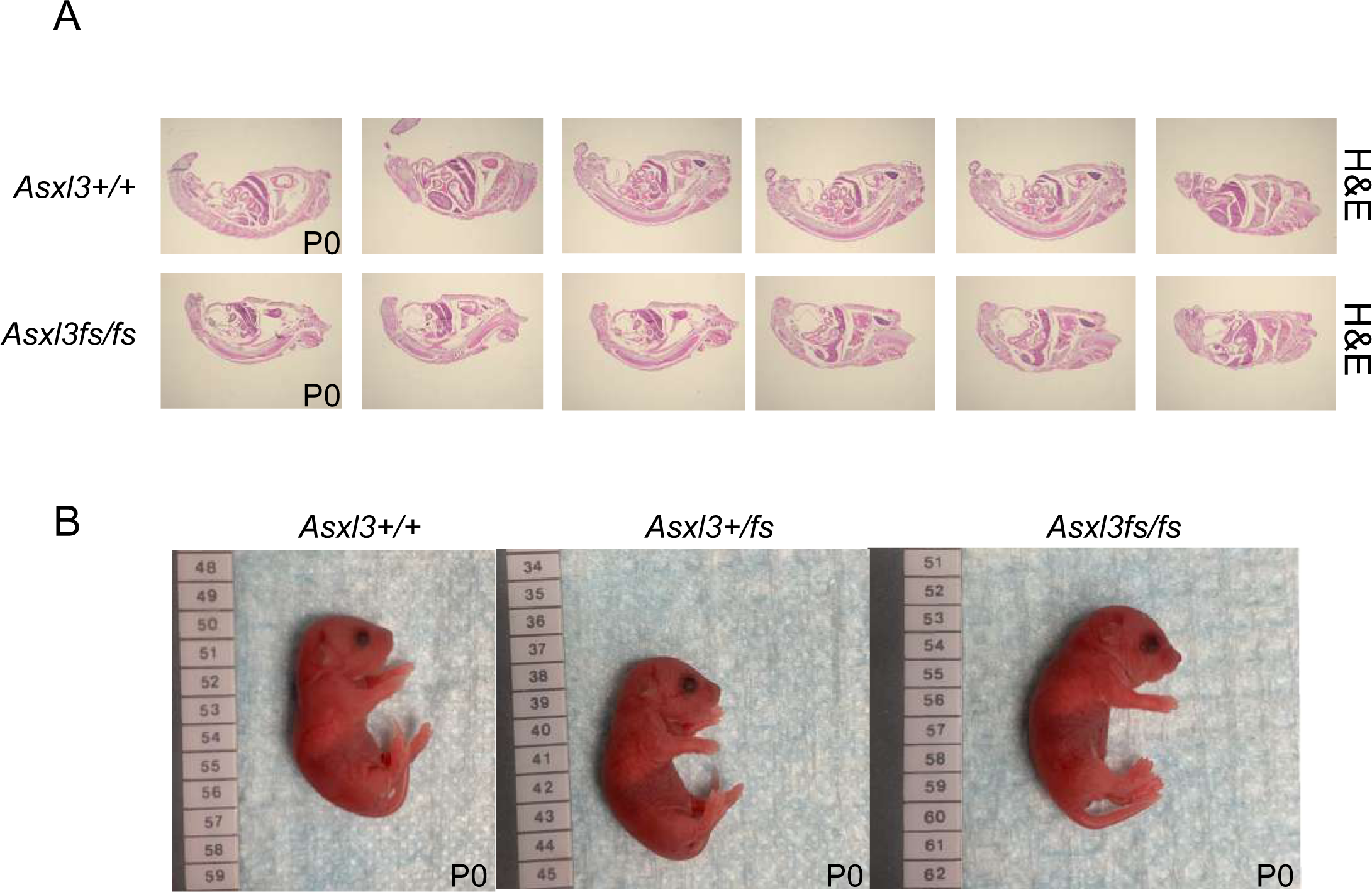
Gross anatomy of Asxl3^fs/fs^ mice. **A,** Representative images of *Asxl3^+/+^* and *Asxl3^fs/fs^* P0 whole body transverse paraffin sections stained with H&E. **B.** Gross appearance of *Asxl3^+/+^*, *Asxl3^+/fs^,* and *Asxl3^fs/fs^* live mice.

**Supplemental Figure 3.**
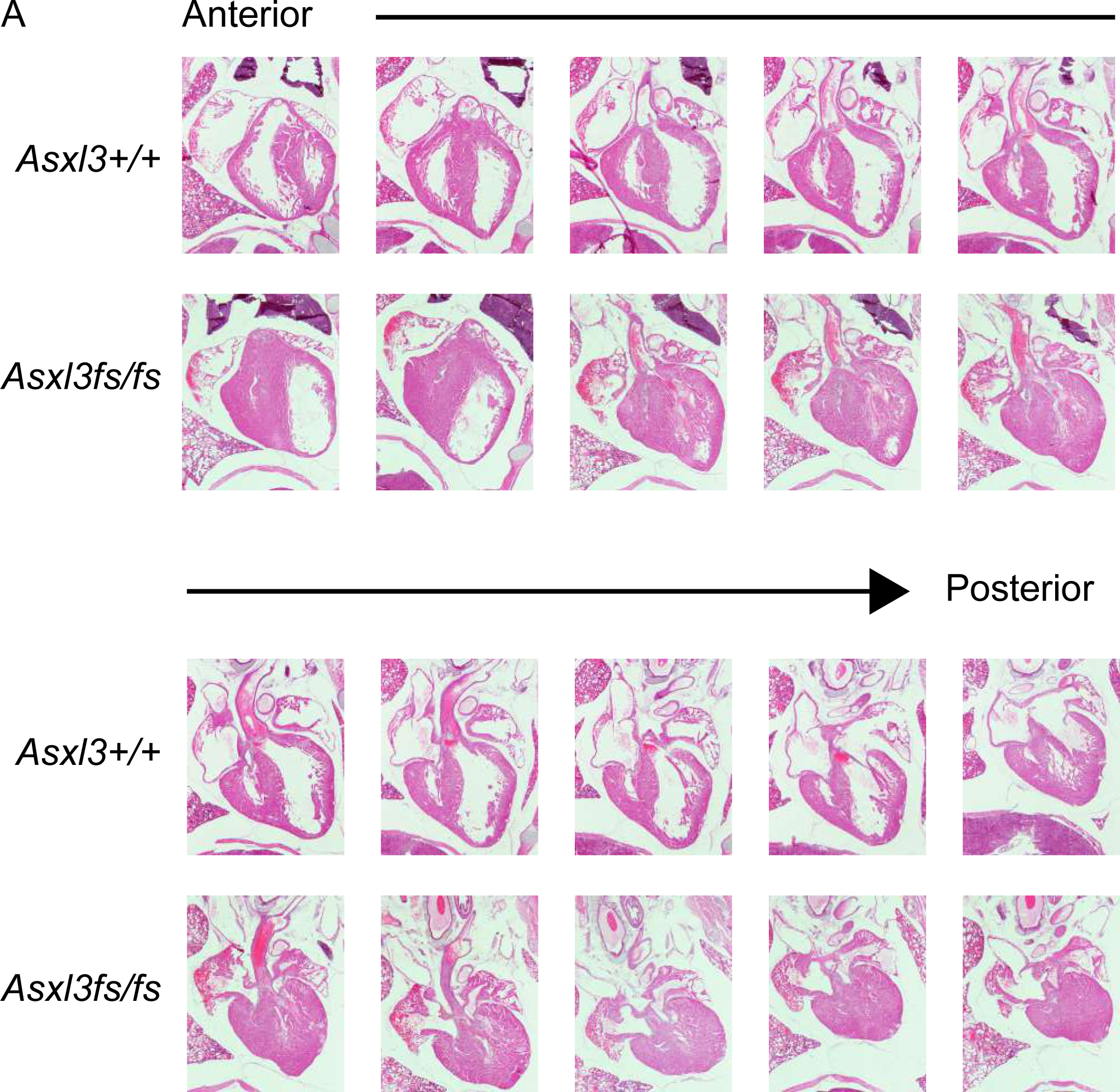
Ventricular hypoplasia phenotype in Asxl3^fs/fs^ mice. **A,** Serial coronal sections of hematoxylin and eosin stained hearts from P0 littermates. Compared with controls, Asxl3^fs/fs^ mice exhibit ventricular hypoplasia throughout the anterior to posterior axis of the heart.

**Supplemental Figure 4.**
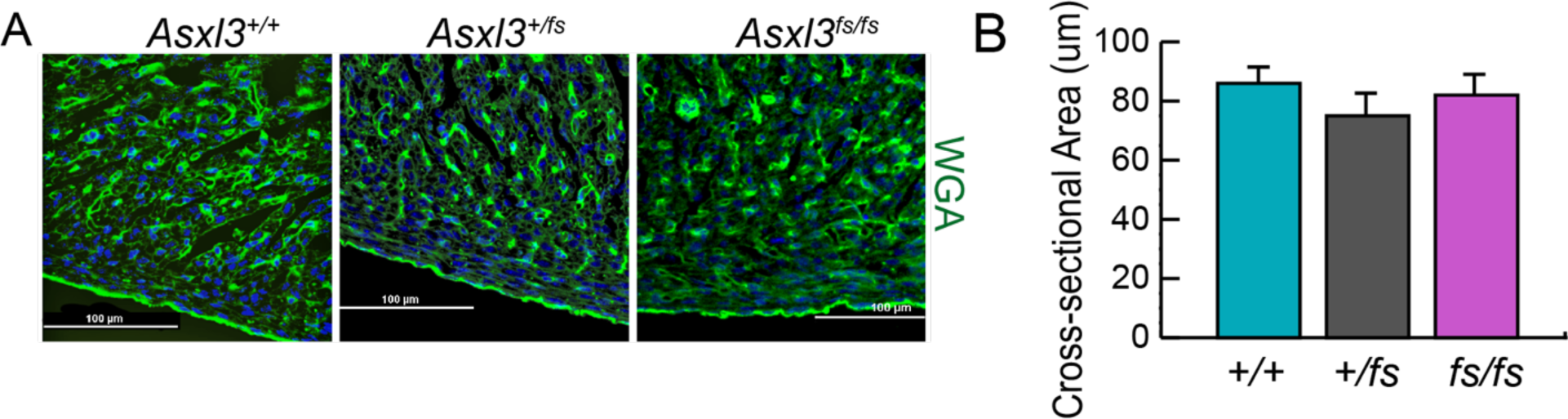
Cross sectional area of cardiomyocytes. **A,** Representative images of *Asxl3^+/+^, Asxl3^+/fs^,* and *Asxl3^fs/fs^* P0 transverse heart sections stained with wheat germ agglutinin (WGA). **B,** Quantification of cardiomyocyte coss-sectional area.

**Supplemental Figure 5.**
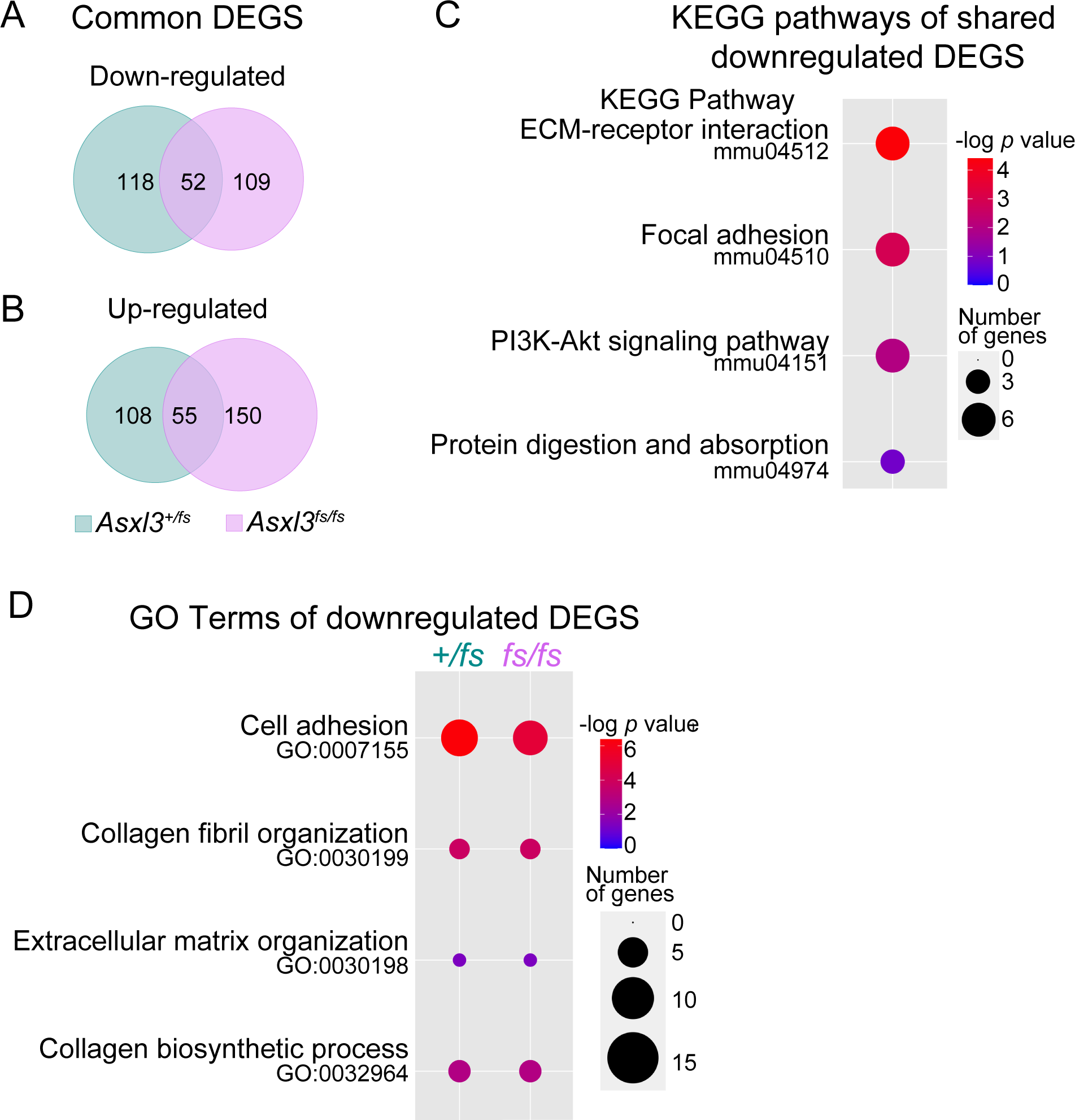
Differentially expressed genes shared by Asxl3^+/fs^ and Asxl3^fs/fs^. Venn diagrams comparing the significantly **A,** downregulated and **B,** upregulated genes (p- value > 0.05 and log fold change > 0.4) for *Asxl3^+/fs^* and *Asxl3^fs/fs^* E18.5 hearts. **C,** KEGG pathway enrichment analysis for shared *Asxl3^+/fs^* and *Asxl3^fs/fs^* downregulated genes. The top 4 KEGG pathways are shown. The size of each dot represents the gene number and the shading represents the -log10 p-value. **D,** Common Gene Ontology terms shared between *Asxl3^+/fs^* and *Asxl3^fs/fs^* downregulated genes. The size of each dot represents the gene number and the shading represents the -log10 p-value.

**Supplemental Figure 6.**
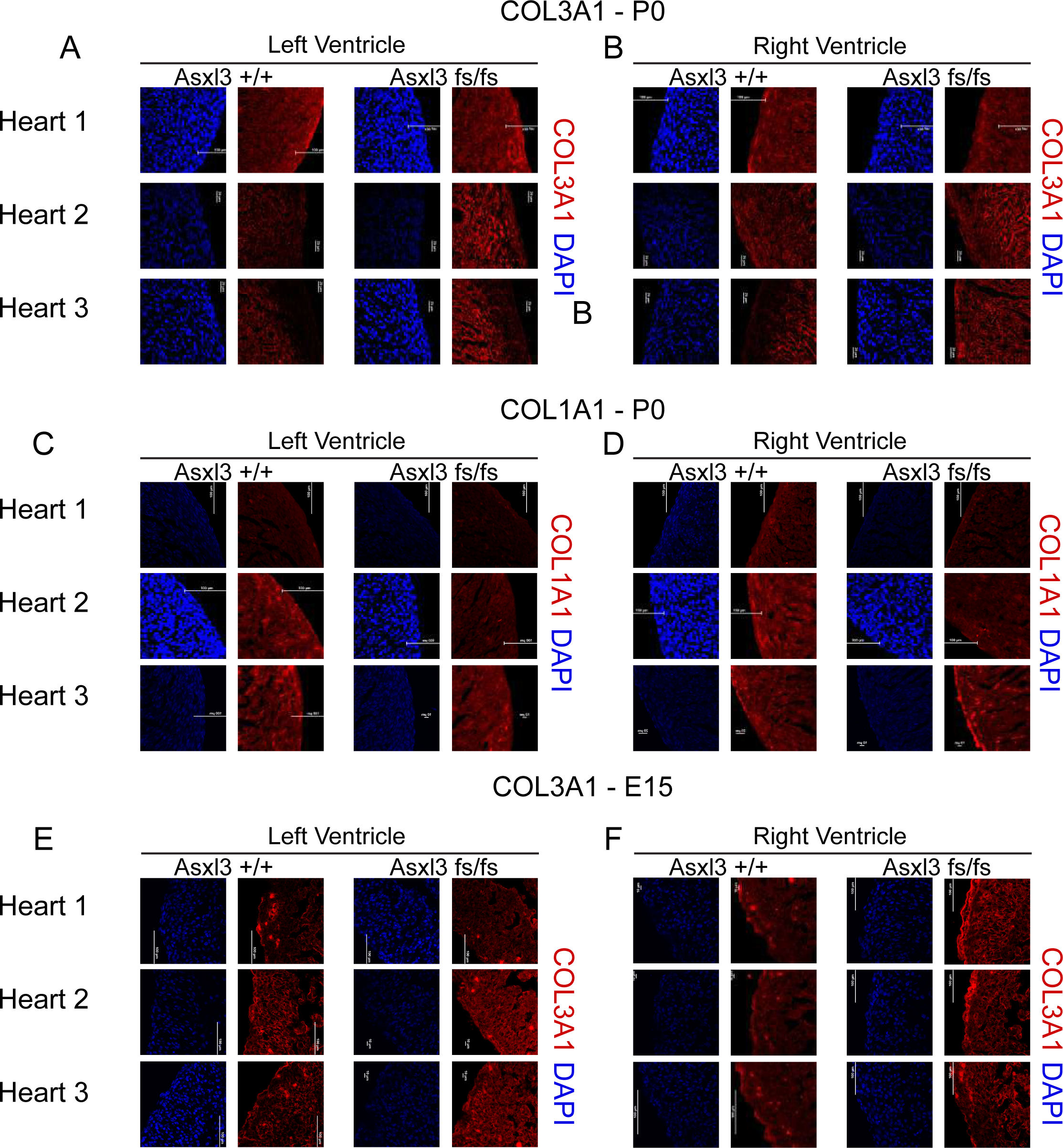
Collagen expression in the developing heart. Representative images of Asxl3^+/+^ and Asxl3^fs/fs^ P0 **A,** left and **B,** right ventricles immunostained with COL3A1. Representative images of Asxl3^+/+^ and Asxl3^fs/fs^ P0 **C,** left and **D,** right ventricles immunostained with COL1A1. Representative images of Asxl3^+/+^ and Asxl3^fs/fs^ E15 **E,** left and **F,** right ventricles immunostained with COL3A1.

**Supplemental Figure 7.**
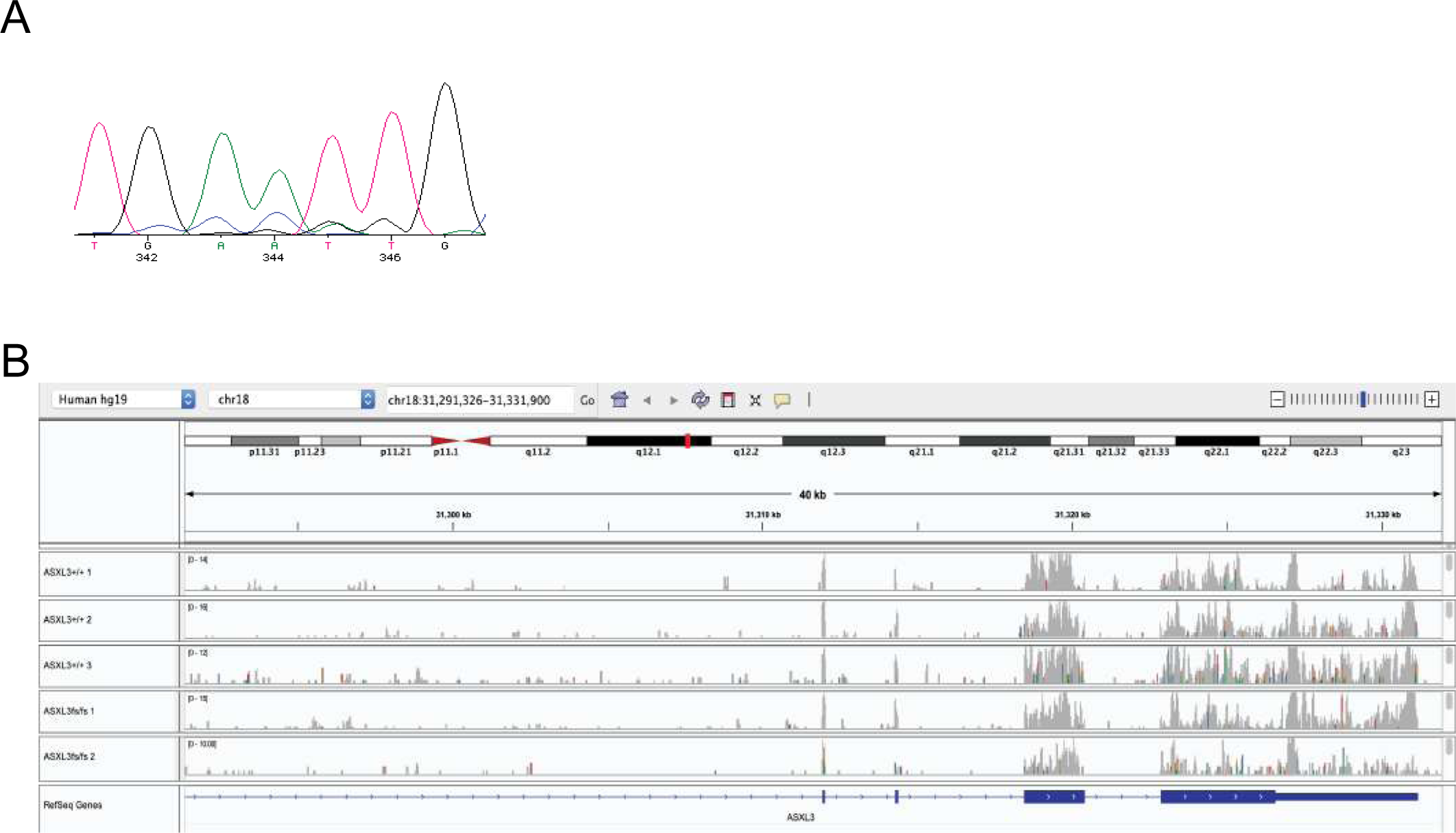
Human embryonic stem cell line carrying compound heterozygous mutation of ASXL3. **A**, Chromatogram from Sanger sequencing of the ASXL3^fs/fs^ cell line identifying a compound heterozygous mutations (c.1393dupT; p.C465LfsX4/c.1390_1393del; p.E464AfsX19) in the ASXL3 gene. **B**, Read density plots focusing on the last 4 exons (Exon 9 through Exon 12) of ASXL3 from the single-cell RNA-Sequencing data of the day 12 cardiac differentiation.

**Supplemental Figure 8.**
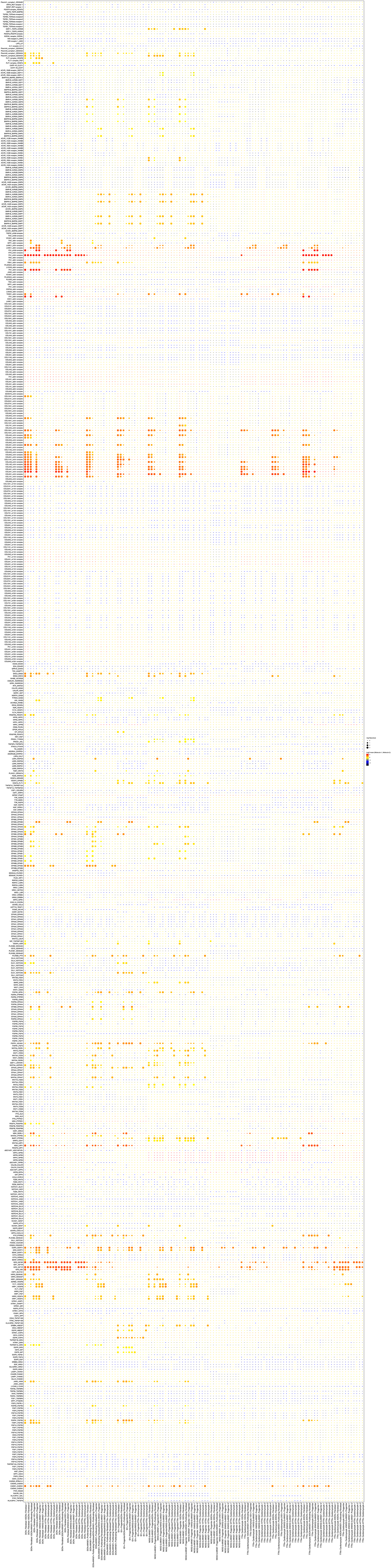
Human embryonic stem cell line carrying compound heterozygous mutation of ASXL3. **A**, Dot plots of all cell-cell communication pathways predicted by CellphoneDB in *ASXL3+/+* cells.

**Supplemental Figure 9.**
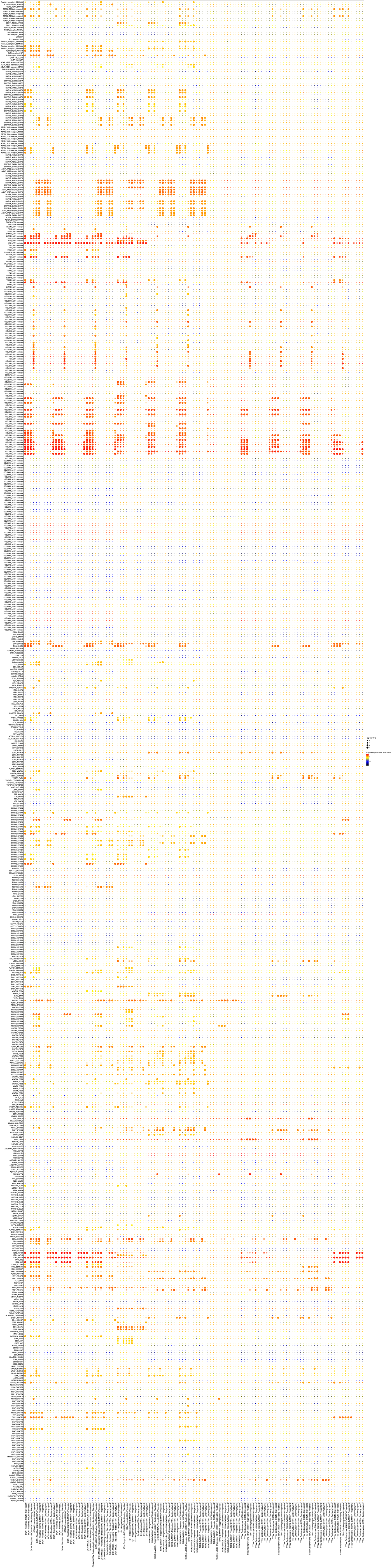
Human embryonic stem cell line carrying compound heterozygous mutation of ASXL3. **A**, Dot plots of all cell-cell communication pathways predicted by CellphoneDB in *ASXL3fs/fs* cells.

**Supplemental Figure 10.**
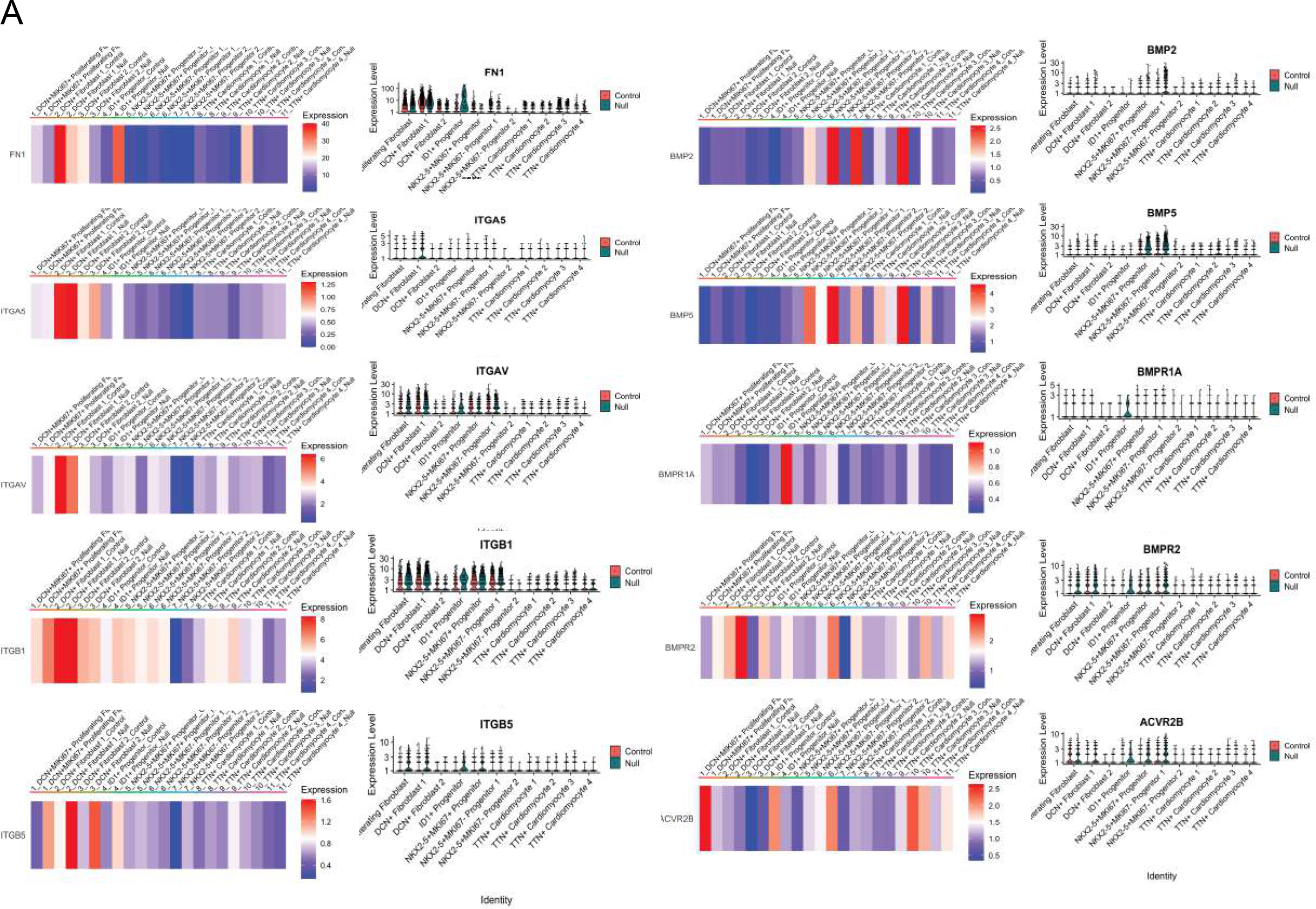
Human embryonic stem cell line carrying compound heterozygous mutation of ASXL3. **A**, Gene expression violin plots and heat maps of the genes shown in Figure 7G in all clusters.

